# Neural Representations of Self-Generated Thought during Think-aloud fMRI

**DOI:** 10.1101/2022.02.15.480505

**Authors:** Hui-Xian Li, Bin Lu, Yu-Wei Wang, Xue-Ying Li, Xiao Chen, Chao-Gan Yan

**Author notes:** **Corresponding author:** Chao-Gan Yan, PhD, CAS Key Laboratory of Behavioral Science, Institute of Psychology, Beijing, China, 16 Lincui Road, Chaoyang District, Beijing 100101, China, E-mail address (C.-G. Yan).

## Abstract

Is the brain at rest during the so-called resting-state? Ongoing experiences in the resting-state vary in unobserved and uncontrolled ways across time, individuals, and populations. However, the role of self-generated thoughts in resting-state fMRI remains largely unexplored. In this study, we collected real-time self-generated thoughts during “resting-state” fMRI scans via the think-aloud method (i.e., think-aloud fMRI), which required participants to report whatever they were currently thinking. We first investigated brain activation patterns during a think-aloud condition and found that significantly activated brain areas included all brain regions required for speech. We then calculated the relationship between divergence in thought content and brain activation during think-aloud and found that divergence in thought content was associated with many brain regions. Finally, we explored the neural representation of self-generated thoughts by performing representational similarity analysis (RSA) at three neural scales: a voxel-wise whole-brain searchlight level; a region-level whole-brain analysis using the Schaefer 400-parcels; at the systems level using the Yeo seven-networks. We found that “resting-state” self-generated thoughts were distributed across a wide range of brain regions, involving all seven Yeo networks. This study highlights the value of considering ongoing experiences during resting-state fMRI, as well as providing preliminary methodological support for think-aloud fMRI.

## 1. Introduction

Resting-state functional magnetic resonance imaging (fMRI) is widely used to study normal cognitive function and to seek biological markers of clinical disease because of its relative simplicity, the ease of accumulating large data, and the ability to reveal intrinsic neural circuitry (Fox and Raichle, 2007; Mulders et al., 2015; van den Heuvel and Hulshoff Pol, 2010). Studies using resting-state fMRI have yielded many findings, however, these can be inconsistent and even contradictory, raising questions about the reliability of the resting-state fMRI method (Buckner et al., 2013; Cole et al., 2010; Kelly et al., 2012). Intra-individual variability in resting-state fMRI is substantial, which reduces its reliability (Zuo and Xing, 2014). The variability of unconstrained thoughts during resting-state likely plays a key role in the variability of outcome measures.

Does “resting state” fMRI data come from a “resting” brain (Finn, 2021)? Can the brain be completely at rest? Recent studies suggest that despite the absence of externally driven cognitive processing during resting-state scanning, individuals still consciously generate much ongoing endogenous cognitive processing, termed self-generated thought (Gorgolewski et al., 2014; Smallwood and Schooler, 2015). Thus, resting-state fMRI data are influenced by many factors, not only non-neural factors such as head motion and scanning conditions (Yan et al., 2009; Yan et al., 2013) but also by the ongoing experience generated by the individual during the resting-state scan and its corresponding neural activity (Buckner et al., 2013; Delamillieure et al., 2010; Gonzalez-Castillo et al., 2019). Although perception and cognition constitute individual ongoing experience at “rest,” researchers rarely account for these contributions, instead applying a single term — intrinsic or spontaneous – to characterize all neuronal activity observed during resting-state fMRI (Gonzalez-Castillo et al., 2021; Kucyi et al., 2018; Morcom and Fletcher, 2007). However, classifying brain activity triggered by ongoing experience as simply intrinsic to the human brain blurs the signals potentially detectable from resting-state fMRI studies. Buckner et al. noted that while resting-state functional connectivity (FC) is not entirely constrained by anatomic connectivity, it is influenced by the mental state of the participant during scanning (Buckner et al., 2013). By comparing dynamic FC analyses between resting and anesthetized states, Barttfeld et al. found that patterns of FC during resting-state also reflected ongoing cognitive processing (Barttfeld et al., 2015); Gonzalez-Castillo et al. found that resting-state dynamic FC was influenced by short periods of spontaneous cognitive-task-like processes (Gonzalez-Castillo et al., 2019). Thus, ongoing experience (self-generated thought) during resting-state fMRI influences the detected signals.

Self-generated thought is a pervasive, complex, and heterogeneous cognitive activity closely associated with various types of mental and psychiatric disorders (Andrews-Hanna et al., 2013; Christoff et al., 2016; Marchetti et al., 2016; Perkins et al., 2015; Wang et al., 2018b). Due to the variable nature of self-generated thoughts (Christoff et al., 2016), the same individual cannot be assumed to produce the same thoughts during two resting-state scans. Thus, failing to control for the effect of resting-state self-generated thoughts can significantly reduce intra-individual consistency across multiple resting-state measures, thus decreasing resting-state fMRI reliability. Another more important issue is that, for intergroup comparison studies, if groups differ in the tendency to produce self-generated thoughts, systematic intergroup differences in their resting state self-generated thoughts will likely be reflected in their resting-state neural activity patterns (Buckner et al., 2013). Notably, psychiatric patients tend to think spontaneously differently from healthy individuals (Andrews-Hanna et al., 2013; Christoff et al., 2016). For example, people with depression have a higher frequency of self-generated thoughts that have more negative content and are more related to the past compared to healthy individuals (Hoffmann et al., 2016). Self-generated thoughts in anxiety disorders are often accompanied by concerns about possible future events (Spinhoven et al., 2015); overactive and assertive self-generated thoughts are associated with an individual’s tendency to mania (Gruber et al., 2008). Moreover, psychotic disorders are characterized by a profound disruption in self-generated thought: frequent abrupt leaps from one topic to another, or rigidity of thought, i.e., repetition or paucity of thought content (Christoff et al., 2016). If the effect of self-generated thought is not considered, the results of intergroup comparisons of resting-state fMRI between people with mental illness and healthy individuals are likely to primarily reflect differences in their self-generated thought, rather than their neural bases. Therefore, for such studies, especially the common comparison studies between patients with mental illness and healthy individuals, systematic differences in resting-state self-generated thought can affect the validity of resting-state fMRI. Therefore, one key to improving the reliability and validity of resting-state fMRI methods is to find ways to accurately measure self-generated thought and control for its effects.

Revealing the ongoing experience of individuals during resting-state fMRI and elucidating their effects on patterns of brain activity during resting-state scans would facilitate the interpretation of resting-state fMRI outcomes and advance the progress of resting-state MRI. However, the role of self-generated thought in resting-state fMRI remains largely unexplored and little progress has been made (Gonzalez-Castillo et al., 2021). Notably, research in the field of mind wandering, which has focused on task-independent cognitive processes, can provide a path for investigating self-generated thoughts in the resting state. Several brain imaging studies have linked the neural basis of mind wandering to the default mode network (DMN). Christoff et al. used the probe-capture experiment to detect when mind wandering occurred, comparing the differences in brain activity in participants when they mind wandered during task completion and when they did not; they found that although the DMN was significantly activated during mind wandering, so was the executive control network (Christoff et al., 2009). Andrews-Hanna et al. manipulated participants’ attention and found that the frequency of spontaneous thoughts was strongly correlated with the strength of activation of the DMN (Andrews-Hanna et al., 2010). Kucyi and Davis used the probe-capture method and dynamic functional connectivity (DFC) analysis and found that DMN DFC was positively correlated with the frequency of mind wandering during task completion (Kucyi and Davis, 2014). These different approaches have implicated the DMN in spontaneous cognitive processing. A meta-analysis of 10 fMRI studies examining spontaneous thought processing found significant activation in several brain regions besides the DMN, including secondary somatosensory cortex, temporopolar, dorsal anterior cingulate cortex, rostrolateral prefrontal cortex, insula and lingual gyrus, emphasizing that the neural basis of self-generated thoughts extends beyond the DMN (Fox et al., 2015). Still, although some progress has been made in identifying the correlates of self-generated thoughts, studies have mainly focused on task contexts, such as sustained attention response tasks or working memory tasks, and have rarely involved examining a task-free or resting state (Fox et al., 2015; Gonzalez-Castillo et al., 2021). Furthermore, investigators typically use the probe-caught method to measure self-generated thoughts (Smallwood and Schooler, 2015), by randomly interrupting participants as they complete a task or take a break and then asking them questions such as “Where was your attention focused before the probe?” This approach provides valuable insights into self-generated thoughts. However, interrupting participants can affect their thought (Seli et al., 2013; Smallwood et al., 2002). Besides, it is an indirect measure that relies on individual introspection (Smallwood and Schooler, 2006, 2015) and has limitations in characterizing the content of self-generated thoughts (Konu et al., 2020; Tusche et al., 2014). More importantly, we cannot observe the flow of thoughts from one state to another in real time.

Although studies of mind wandering can provide insights into self-generated thoughts at rest, studies of mind wandering were usually based on a task context, which cannot be equated with the experience of being at rest. Current studies of resting-state ongoing experiences primarily use the retrospective approach in which participants are asked to assess their thoughts generated throughout the resting-state along various dimensions after completing the resting-state scan (Delamillieure et al., 2010; Diaz et al., 2014; Diaz et al., 2013; Gonzalez-Castillo et al., 2019; Gorgolewski et al., 2014; Karapanagiotidis et al., 2020; Vatansever et al., 2020; Wang et al., 2018a). Delamillieure designed the Resting State Questionnaire, which classifies experiences during resting-state scans into five categories. The investigators administered it to 180 individuals and found that most participants showed a tendency to have some kind of content as the dominant mental activity, with the largest proportion of individuals reporting predominantly visual mental imagery, followed by inner language (Delamillieure et al., 2010). Gorgolewski et al. used retrospective questionnaires to measure individuals’ self-generated thoughts during resting-state scanning and examined the association of the content and the form of thoughts with resting-state metrics. Results showed that individuals who reported more imagery exhibited greater fractional amplitude of low frequency fluctuations (fALFF) in the perigenual cingulate cortex, and individuals who reported more future thoughts exhibited lower regional homogeneity (ReHo) in the lateral occipital cortex (Gorgolewski et al., 2014). Similarly, Wang et al. tested the correlation of certain brain states (functional connectivity, FC) with retrospective self-reports of experience during the resting-state scan and found variations along four dimensions of thought content. For example, visual experiences were associated with stronger FC between visual and other networks (Wang et al., 2018a). Although the retrospective method can inform on the relationship between general characteristics of resting-state ongoing experience and neural activity, this method lacks temporal precision and lacks individual instances that reveal ongoing mental activity, i.e., it cannot examine real-time experience. Additionally, the retrospective method aggregates information over a long period, blurring details of content, shape, and quality of a particular event experienced in a stationary progression occurring at any given moment (i.e., the temporal blurring effect) (Gonzalez-Castillo et al., 2021). Finally, the retrospective method relies heavily on memory and has a strong lag and memory bias.

How do we make a direct connection between the brain and the mind? The “Spatiotemporal Neuroscience” proposed by Northoff et al. (Northoff et al., 2020) stated that the temporo-spatial dynamics provided the “common currency” that connected neuronal and mental activity. They emphasized that the spontaneous activity of the brain constructs its “inner time and space” and that the different ways in which spontaneous activity constructs its “inner time and space” manifest as different mental features (Northoff and Zilio, 2022). Spatiotemporal Neuroscience conceptualizes the brain in dynamic rather than functional terms, pointing out that we need to shift the focus from task-evoked activity to the spontaneous activity of the brain, i.e., the resting state, emphasizing the construction of the brain’s internal time and space (Northoff et al., 2020). In terms of self-generated thoughts, the dynamic framework proposed by Christoff et al. has profound implications for our understanding of self-generated thoughts. It highlights the importance of exploring how mental states change over time (Christoff et al., 2016). Researchers using electroencephalogram (EEG) techniques have demonstrated that the dynamics of the mind can be tracked by neurodynamics (Hua et al., 2022). Besides, Rostami and his colleagues found that the thought dynamics of patients with depression were altered compared to healthy control (Rostami et al., 2022). They both used the probe-capture method to focus on the internal-external thought dynamics of self-generated thought. Our previous behavioral study showed that participants could express their thoughts and validated the feasibility and reliability of the think-aloud method (Li et al., 2021). Specifically, we found that the think-aloud method did not significantly change the frequency or content characteristics of self-generated thoughts; moreover, its test-retest reliability was high. Additionally, Raffaelli and Andrews-Hanna’s recent work also showed that the think-aloud paradigm could reveal the content, dynamics, and conceptual scope of resting-state thoughts (Raffaelli et al., 2021). The major benefit of think-aloud over probe-caught and retrospective methods is that it is a continuous measure of naturalistic thought, and the data truly offers a sense of thought transitions over time. Therefore, the present study sought to collect real-time, immediate resting-state self-generated thoughts with the goal of examining how an individual’s brain activity patterns were influenced by his or her self-generated thoughts, and to clarify the direct form of brain representations of self-generated thoughts and the feasibility of the think-aloud fMRI method to study ongoing experiences. Specifically, we used an immediate think-aloud method to collect self-generated thoughts during a “resting-state” fMRI scan, i.e., the think-aloud fMRI method. This method required participants to report any thoughts or images that occurred in their minds during a 10-minute think-aloud fMRI scan, which we note, is not a real “resting-state” scan. We used MRI to synchronize fMRI data throughout the process and recorded participants’ verbal reports with an MRI-compatible adaptive noise canceling microphone.

Thoughts in the resting-state arise and proceed in a relatively free, unconstrained fashion (Mildner and Tamir, 2019). When a person is at rest, his or her brain is often filled with a variety of experiences, meaning that the individual experiences a wealth of mental activity during a resting-state fMRI scan, including visual mental imagery, inner language, auditory mental imagery, somatosensory awareness, inner musical experience and mental processing of numbers (Delamillieure et al., 2010; Diaz et al., 2014). Thus, a wide range of brain areas can be involved. For example, visual mental imagery corresponds to visual areas, somatosensory awareness corresponds to sensorimotor areas, and internal language corresponds to language areas. Furthermore, self-generated thoughts involve a great deal of episodic memory; at least 50% of self-generated thoughts are about the future and the past (Klinger and Cox, 1987; Smallwood and Schooler, 2015; Wamsley, 2019). Researchers have proposed that self-generated thoughts and memory processes have much in common and emphasized that semantic memory provides the scaffolding for episodic memory to shape the content of thought (Mildner and Tamir, 2019). Huth et al. has systematically mapped semantic selectivity across the cortex and found that the semantic system was organized into intricate patterns relatively symmetrically in the two cerebral hemispheres (Huth et al., 2016). Moreover, they found that different areas of the brain responded to different kinds of words and concepts. But this was not just one word linked to one location, as a single word activated a whole range of diverse brain regions. Given the complexity and rich semantic features of resting-state experience, we believed that the brain representation of experiences during resting-state would be intricate and involve a wide range of brain regions across all networks, rather than being restricted to a single or a few networks.

In the present study, we first explored brain activation patterns when participants reported their thoughts. To further illustrate a relationship between the activation of verbally reported self-generated thoughts and thought content, we calculated correlations between brain activation patterns and divergence in thought content using an indicator that was shown to have good reliability and validity (Li et al., 2021). Finally, we tried to detect direct neural representations of self-generated thoughts in the “resting-state” scan through representational similarity analysis (RSA), which can effectively clarify relationships between different data modalities (Kriegeskorte and Kievit, 2013; Popal et al., 2019). Specifically, for participants’ verbally reported thoughts, we transcribed and labeled each self-generated thought episode based on topic similarity and then used NLP to convert each self-generated thought episode into a fixed-length vector for quantitative analysis. This approach is effective for studying self-generated thoughts in the resting-state (Li et al., 2021). For the fMRI data, we calculated average BOLD (blood oxygenation level-dependent) values to obtain brain activity patterns of each self-generated thought episode. Based on the meta-analysis of spontaneous cognitive processing (Fox et al., 2015) and the diverse characteristics of resting-state experiences (Delamillieure et al., 2010), we hypothesized that self-generated thoughts during resting-state fMRI would involve a wide range of brain areas, and not only be limited to the DMN and the cognitive control network but would also involve attentional and sensorimotor networks, among others. Therefore, we examined the brain representation of self-generated thoughts using RSA at three whole-brain neural levels: a voxel-level, using whole-brain searchlight analysis; a region-level, using the Schaefer 400-parcels (Schaefer et al., 2018); and a systems-level, using the seven Yeo networks (Yeo et al., 2011)(Fig. 1).

**Fig. 1.**
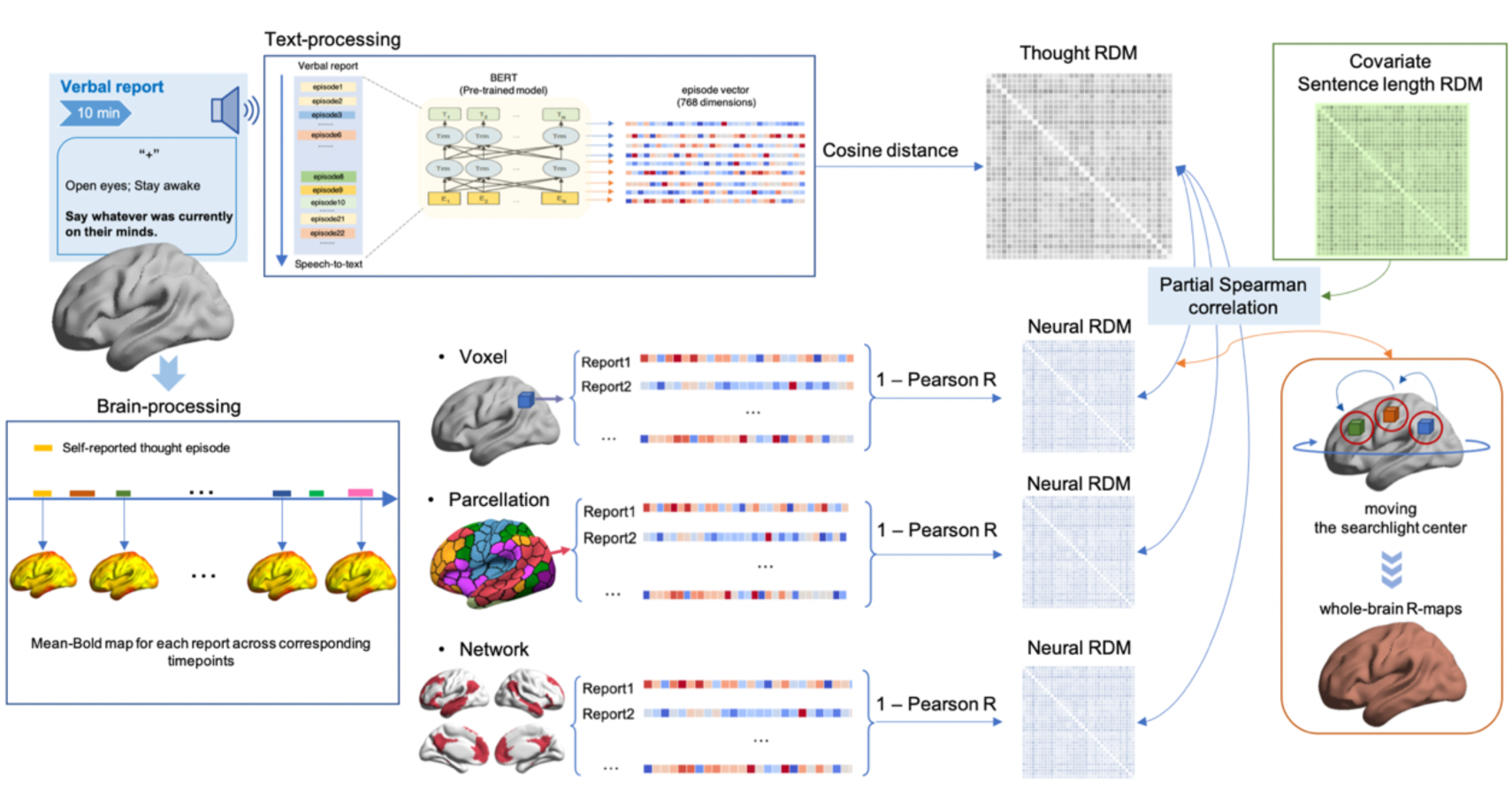
The RSA schematic diagram. The thought RDM was constructed by calculating the cosine distance of the BERT vector between two thought episodes; the sentence length RDM was constructed by calculating the absolute value of word count of two thought episodes. Brain activity maps for each reported thought episode were obtained using each thought episode as a condition. RDMs for the three neural levels were constructed by calculating 1 minus the Pearson correlation between the BOLD values of all voxels contained within each thought episode computational unit (voxels, parcels, and networks). The partial Spearman correlation was used to calculate the correlation between the thought RDM and the neural RDM, with the sentence length RDM as a covariate.

## 2. Materials and methods

### 2.1. Participants

One hundred and one healthy adult participants recruited by internet advertisements completed the think-aloud fMRI experiment. People who reported any MRI contraindications, psychiatric or neurological disorders, use of psychotropic medications, or any history of substance or alcohol abuse were excluded. Two participants were excluded because the device was not recording. Two subjects who did not report any thoughts were excluded. Three participants whose voices were too low to be accurately transcribed were excluded. Five participants whose verbal reports contained fewer than five thoughts were excluded. Three participants were excluded because their head motion exceeded the criterion (mean Framewise Displacement (FD) Jenkinson > 0.2 mm) (Jenkinson et al., 2002). Therefore, the sample size for the final data analysis was 86 (45 females; mean age = 22.1 ± 2.7 years). All participants gave informed written consent. All experiments were approved by the Institutional Review Board of the Institute of Psychology, Chinese Academy of Sciences.

### 2.2. Experimental design

Several days before scanning (no more than one week apart), participants were interviewed and informed of the purpose of the study and practiced producing think-aloud verbal reports outside the scanner. We briefly introduced the concept of self-generated thoughts and asked a few questions to deepen their understanding of this phenomenon. All introduction and interview information were the same as previously reported (Li et al., 2021). Briefly, we communicated the following. “When we are resting, the brain doesn’t completely rest. Our minds are often filled with thoughts or images, commonly known as ‘daydreaming’, ‘drifting’ and so on. In addition to the resting state, when we are in states of working, studying, listening to lectures, etc., our minds are often filled with thoughts or images that are not related to the activity we are doing, a condition we usually refer to as ‘wandering’. Moreover, when we are sleeping, our brain also produces a lot of thoughts and images, which is called ‘dreaming’. Our brains often produce a series of changeable thinking activities without a specific destination. And most of these thinking activities are endogenous, separate from the external perceptual process. In psychology, this phenomenon is called ‘mind wandering’, or ‘spontaneous thinking’.” We also informed participants that the reporting periods would be audio-recorded throughout although we would not listen to their reports concurrently; that the audio-recording file names would not contain any personally identifiable information; and that we would only use the text for scientific research and would not disclose any of their personal information. Subsequently, participants performed a 5-minute verbal report exercise without audio recording to adapt to the think-aloud paradigm. Afterward, participants completed a 10-minute formal verbal report exercise with audio recording.

One hundred and one participants finished a free verbal report stage of the think-aloud fMRI method. Participants were asked to look at a fixation cross on the screen with their eyes open and to stay awake. This was a classic resting-state scan with the difference that any thoughts and images needed to be reported directly when the participant became aware of them, without regard to grammar. We emphasized that participants were not required to think deliberately and to maintain a natural resting state as much as possible. The purpose of the exercise was to achieve self-talk as much as possible, i.e., to keep up with their thinking as much as is possible verbally. In addition, at the end of the scan participants rated their thought content (temporal, social, and emotional valence) and form (words and images) on a scale from 1 to 9 points (Gorgolewski et al., 2014) (Table S1). We used the FOMRI-III^TM^ (Fiber Optic Microphone for Functional MRI, https://www.optoacoustics.com/medical/fomri-iii/features), which is an advanced adaptive noise canceling microphone used in MRI environments to record participants’ verbal reports.

### 2.3. MRI data acquisition

MRI data were acquired on a GE MR750 3.0T scanner with 8-channel head coil at MRI Research Center, Institute of Psychology, Chinese Academy of Sciences. The 3D T1-weighted images were acquired using 3D-SPGR pulse sequence (192 sagittal slices, repetition time (TR) = 6.65 ms, echo time (TE) = 2.93 ms, flip angle (FA) = 12 degrees, inversion time (IT) = 450 ms, field of view (FOV) = 256 × 256 mm^2^, matrix size = 256 × 256, slice thickness = 1 mm, voxel size = 1 × 1 × 1 mm). Functional data were acquired with an echo-planar imaging (EPI) sequence (37 axial slices, TR = 2000 ms, TE = 30 ms, FA = 90 degrees, FOV = 224 × 224 mm^2^, matrix size = 64 × 64, slice thickness = 3.5 mm, voxel size = 3.5 × 3.5 × 3.5 mm).

As speaking-induced head motion might be a concern for the think-aloud fMRI, we adopted some strategies to control participants’ head motion. 1) A previous study reported that applying a medical tape from one side of the head coil to the other side over the participant’s forehead significantly reduced head motion (Krause et al., 2019), so we applied this strategy for almost half of our participants (n = 50). However, we didn’t detect a significant head motion difference between the participants who were scanned with or without the medical tape (t = 0.539, p = 0.591). 2) Padding sponges were used to fill the gap between the left and right sides of the MRI head coil and participants’ head. 3) We requested that participants move their mouths as steadily as possible while speaking and emphasized the importance of controlling head motion. 4) We monitored participants’ head motion in real-time during scanning. When participants exceeded the real time maximum head motion criteria (3 mm or 3 degrees on any axis) during the scan, we aborted the scan to restart a new scan with additional emphasis on the need to control head motion. Despite all the strategies we applied, 3 participants still had mean FD_Jenkinson greater than 0.2 mm after postprocessing. As they exceeded the head motion criterion, we excluded these participants from analyses.

### 2.4. Verbal report data preprocessing

As in our previous study (Li et al., 2021), we converted verbal reports into text with the speech-to-text platform iFLYTEK (https://www.iflyrec.com) with manual supervision. Various stop words such as “um” and “ah” were discarded using iFLYTEK. We manually labeled thought switches according to chronological order and the similarity of the topics and calculated the number of self-generated thought episodes. Specifically, cuts were made when different topics appeared in chronological order. For example, “In the evening, I met someone to go for a walk” followed by “In a minute I’m going to call my sister” was marked as a thought switch between two different self-generated thought episodes, as the two represented different topics. Conversely, “In the evening, I met someone to go for a walk” followed by, “Told him to go to a nearby park” was not marked as a thought switch because it was the same topic. Text proofreading and thought switch markers were done independently by a psychology student at three different times to ensure the correctness of the transcriptions, in addition to consulting with others when different markers were encountered. Our previous study showed that the test-retest reliability of labeled thought transition frequencies was good (the intraclass correlation coefficient, ICC > 0.8) (Li et al., 2021). Finally, we confirmed the start and end time of each reported self-generated thought episode in preparation for brain imaging data analyses. Then, we mapped the sentences contained in each self-generated thought episode into a 768-dimensional fixed-length vector representation using the BERT model (Devlin et al., 2019) and bert-as-service (https://github.com/hanxiao/bert-as-service). The pre-trained BERT model was Chinese_L-12_H-768_A-12 provided by Google. BERT is a deep learning model that generates word vector representations in sentences as well as sentence vector representations that can be used for both word-level natural language processing tasks and sentence-level tasks. In our study, we used sentence-level representations.

### 2.5. fMRI data preprocessing

We used the DPABI (Data Processing & Analysis for Brain Imaging, http://rfmri.org/dpabi) (Yan et al., 2016) toolbox which is based on SPM (Statistical Parametric Mapping, http://www.fil.ion.ucl.ac.uk/spm) to preprocess the fMRI data. The preprocessing was as follows: The initial 10 volumes (20 s) were removed. Slice-timing correction (Sladky et al., 2011) was performed with all volume slices corrected for different signal acquisition times by shifting the signal measured in each slice relative to the acquisition of the slice at the mid-point of each repetition time. Then, realignment was performed using a six-parameter (rigid body) linear transformation with a two-pass procedure (registered to the first image and then registered to the mean of the images after the first realignment). After realignment, individual T1-weighted MPRAGE images were co-registered to the mean functional image using a 6 degree-of-freedom linear transformation without re-sampling and then segmented into gray matter, white matter (WM), and cerebrospinal fluid (CSF). The new segmentation method of SPM and the Diffeomorphic Anatomical Registration Through Exponentiated Lie algebra (DARTEL) tool (Ashburner, 2007) were used to segment and calculate the information of spatial transformation. Then, functional images were normalized to MNI space by DARTEL, with the voxel size resampled to 3 × 3 × 3 mm. Finally, all functional images were smoothed with a 4 mm FWHM Gaussian kernel. The preprocessing decisions were similar to the study by Chen et al. (Chen et al., 2017). We did not conduct global signal regression (GSR) because it is still controversial (Murphy and Fox, 2017).

### 2.6. Analysis of activation results with think-aloud fMRI

SPM12 was used for individual-level analyses; a general linear model (GLM) was estimated for each voxel. The design matrix of the free verbal report stage consists of each report and the six head motion parameters computed at the realignment stage. Here, all reported self-generated thought episodes were designed as a single condition. The regressors were convolved with the SPM canonical hemodynamic response function. The high-pass filter was set at 128 s. After GLM estimation, we obtained the beta-map of reporting in the free verbal report stage. Group-level analyses were assessed using whole-brain con-weight images. Multiple comparisons correction was performed using the Gaussian Random Field (GRF): voxel-level p < 0.001, cluster-level p < 0.05.

### 2.7. Divergence in thought content

Like our previous study (Li et al., 2021), we first divided each 10-minute stage into two five-minute segments. Then, we calculated the sum vector of all self-generated thought episode vectors (768-dimensional token vectors from the final layer of BERT) to represent the content of the segment. Finally, we computed the inverse cosine value between two sum vectors to detect the overall divergence of thought content between the two segments (Fig. 3). This indicator reflects variability in the content of ideas over time. It indicates the similarity of text content between two-time segments; the smaller the value, the more similar is the thought content of the two-time segments. Then, we conducted a correlation analysis between the divergence indicator of reported thought content and the beta-map of the free verbal report stage, for which all reported thought events were organized as a single condition in the design matrix. In addition, we performed a parcel-level analysis using Schaefer 400-parcels. We averaged the activation values of all voxels within a parcel. The 400 p-values were submitted to FDR correction (q = 0.05). This analysis was performed to clarify the brain regions associated with the thought contents.

### 2.8. Representational similarity analysis on three neural scales

We obtained the content of participants’ self-generated thought episodes to construct their thought content representational dissimilarity matrix (RDM) using NLP for quantitative analysis. We also obtained each individual’s BOLD fMRI responses for each self-generated thought episode. Then, neural RDMs were constructed for each individual using the thought episode pairwise correlations of BOLD response patterns at three different scales: a voxel-wise analysis with a whole-brain searchlight method; a region-level analysis with the Schaefer 400-parcels; at a systems level with the seven Yeo networks and language subsystems (Fig. 1).

#### 2.8.1. Building RDM based on the content of thought episodes

We obtained each participant’s reported thought RDM based on the BERT vectors of each self-generated thought episode. Specifically, the semantic distance was measured as the cosine distance between BERT vectors of each self-generated thought episode pair. Furthermore, we calculated the number of words reported for each self-generated thought episode to assess differences in sentence length. We observed a total of 1938 thought episodes (Mean: 37.600; SD: 30.316). The distribution of sentence lengths is shown in Fig. S3 of the supplementary material. Sentence lengths of self-generated thought episodes differed. Therefore, to avoid our main results reflecting only differences in thought sentence length, the sentence length RDM was used as a covariate in our main RSA analysis to control for the effects of episode length. Each participant’s sentence length RDM was constructed by calculating the absolute difference in sentence lengths between each self-generated thought episode pair.

#### 2.8.2. Obtaining whole-brain activity patterns for each reported thought episode

Participants differed in the number of self-generated thought episodes (Mean = 22, SD = 10.09, Fig. 3C). By reference to the way brain imaging data were processed when participants watched (movie-viewing) and recalled naturalistic stimuli (spoken-recall) (Chen et al., 2017; Zadbood et al., 2017), we calculated the average of the BOLD values for the timepoints (TRs) contained in each episode to provide a single brain response pattern for each episode. The RDM for different levels of brain activity was then calculated as follows.

#### 2.8.3. Voxel-wise whole-brain searchlight analysis

We performed a searchlight analysis (Kriegeskorte and Kievit, 2013) to determine the similarity of activity patterns across voxels between different thought episodes by using NeuroRA (Lu and Ku, 2020), a python toolbox of representational analysis for multi-modal neural data. Parameters were set as follows: ksize = [5, 5, 5]([k_x_, k_y_, k_z_]), with Ksize referring to the number of voxels along the corresponding axes. Strides = [1,1,1] ([s_x_, s_y_, s_z_]]), with Strides indicating how far the calculation unit is moved before another computation is made. Specifically, for each voxel in each participant, we extracted the BOLD values of neighboring voxels forming a cubic region of interest (calculation unit). Then, we constructed the neural RDM by calculating 1 minus the spatial Pearson correlation between voxel-wise BOLD values of different thought episodes within this calculation unit. The neural RDM at a given voxel was compared with the thought RDM using partial Spearman correlation, in which the sentence length RDM was covaried. By moving the calculation unit center throughout the cortex, we obtained whole-brain r-maps and performed Fisher’s r-z transformation. As each voxel can belong to multiple calculation units (Lu and Ku, 2020), the final result for a voxel was the average of all kernels containing that voxel (Fig. 1). Finally, we conducted a group-level one-sample T-test. Multiple comparisons correction was performed using the Gaussian Random Field (GRF): voxel-level p < 0.001, cluster-level p < 0.05 using the grey matter mask in DPABI. Finally, voxel-wise whole-brain searchlight analysis results were matched to the seven Yeo networks (Yeo et al., 2011) for interpretation. Specifically, we calculated the distribution of significant voxels in the Yeo networks and the ratio of significant voxels in each network.

#### 2.8.4. Region-level RSA

By using the Schaefer 400-parcels (Schaefer et al., 2018), region-level whole-brain RSA was performed according to the following computational procedure (Fig. 1): 1) the Schaefer 400-parcels atlas was resampled to 3 × 3 × 3 mm (nearest neighbor interpolation) to match the resolution of our preprocessed data. 2) the BOLD value was extracted per parcel for each thought episode; 3) the neural RDM was computed as the 1 minus spatial Pearson correlation between BOLD values of different thought episodes for each parcel; 4) the partial Spearman correlation was computed between each parcel’s RDM and thought RDM, controlling for the sentence length RDM as a covariate; and 5) Fisher’s r-to-z algorithm was applied to conduct a group-level one-sample T-test. Parcel-level p-values were submitted to Bonferroni correction (p = 0.05/400). Finally, significant parcels were categorized per the seven Yeo networks and the proportion of significant parcels in each network was calculated.

#### 2.8.5. Systems-level RSA

In systems-level analysis, Yeo’s seven networks were used as masks for RSA (Fig. 1). The analysis process was like the above-mentioned region-based approach but used network masks instead of parcels.

### 2.9. Validation analysis

We conducted supplemental analyses to confirm the validity of our results. 1) We calculated the correlation between the individual mean FD_Jenkinson index during the free verbal report stage and her/his beta-map to see if head motion influenced results (n = 86). 2) We randomly permuted participants’ thought RDMs and recalculated the RSA analysis to determine whether our results reflected random noise. 3) For the fMRI data, we also used the GLM-style analysis to obtain the brain activity pattern of each self-generated thought episode. Each participant had a different number of self-generated thought episodes. SPM12 was used for the first-level analysis to obtain whole-brain activity patterns for each reported thought episode based on the individual level. Specifically, for each self-generated thought episode reported by each participant, a general linear model was estimated for each voxel. The design matrix included the onsets of each self-generated thought episode and six head motion parameters. The high-pass filter was set at 128 s. Every self-generated thought episode was considered a separate regressor. The regressors were convolved with the SPM canonical hemodynamic response function. After model estimation, we obtained the whole-brain beta-weight image for each self-generated thought episode for each participant. Then, we recalculated the RSA at three neural levels using the beta-map of each episode. 4) We calculated another divergence indicator in thought content (Fig. S10). Specifically, we first computed the average vector of self-generated thought episodes (10-minute) and then calculated the difference between each self-generated thought episode and this average vector using the inverse cosine value. Finally, we calculated the coefficients of variation (CV) of these differences to characterize the magnitude of the change/oscillation of the thought content over time throughout the 10-minute stage. We named this indicator a thought content fluctuation indicator. Both the fluctuation indicator and the divergence indicator reflect changes in thought content. The divergence indicator reflects the difference in thought content before and after larger scale times, while the fluctuation indicator reflects the fluctuation in thought content over time. We calculated the correlation between this indicator and the beta-map of the free verbal report stage at the voxel and parcels level, which was the same as the divergence indicator analysis.

### 2.10 Data/code availability statement

All codes are available on git-hub (https://github.com/Chaogan-Yan/PaperScripts/tree/master/LiHX_2022). Data have been shared online (http://rfmri.org/ThinkAloudfMRIData). Of note, the thought vectors were shared rather than the original voice recordings to protect participants’ privacy.

## 3. Results

### 3.1 Brain activation patterns during free verbal reports

We first explored brain activation patterns during reporting, on the one hand, to clarify whether language reporting activated language areas (indicating at least that the fMRI task was successful) and, on the other hand, to map the language areas where we should be cautious in interpreting their relationship with self-generated thoughts. As expected, the think-aloud process significantly activated language-related areas that were consistent with the brain regions required for “speech” (Fig. 2B). It is worth noting that besides the significant activation of speech-related brain areas, many brain areas were deactivated. Although deactivation is not the focus of the current study, its meaning and mechanism need further exploration in the future.

**Fig. 2.**
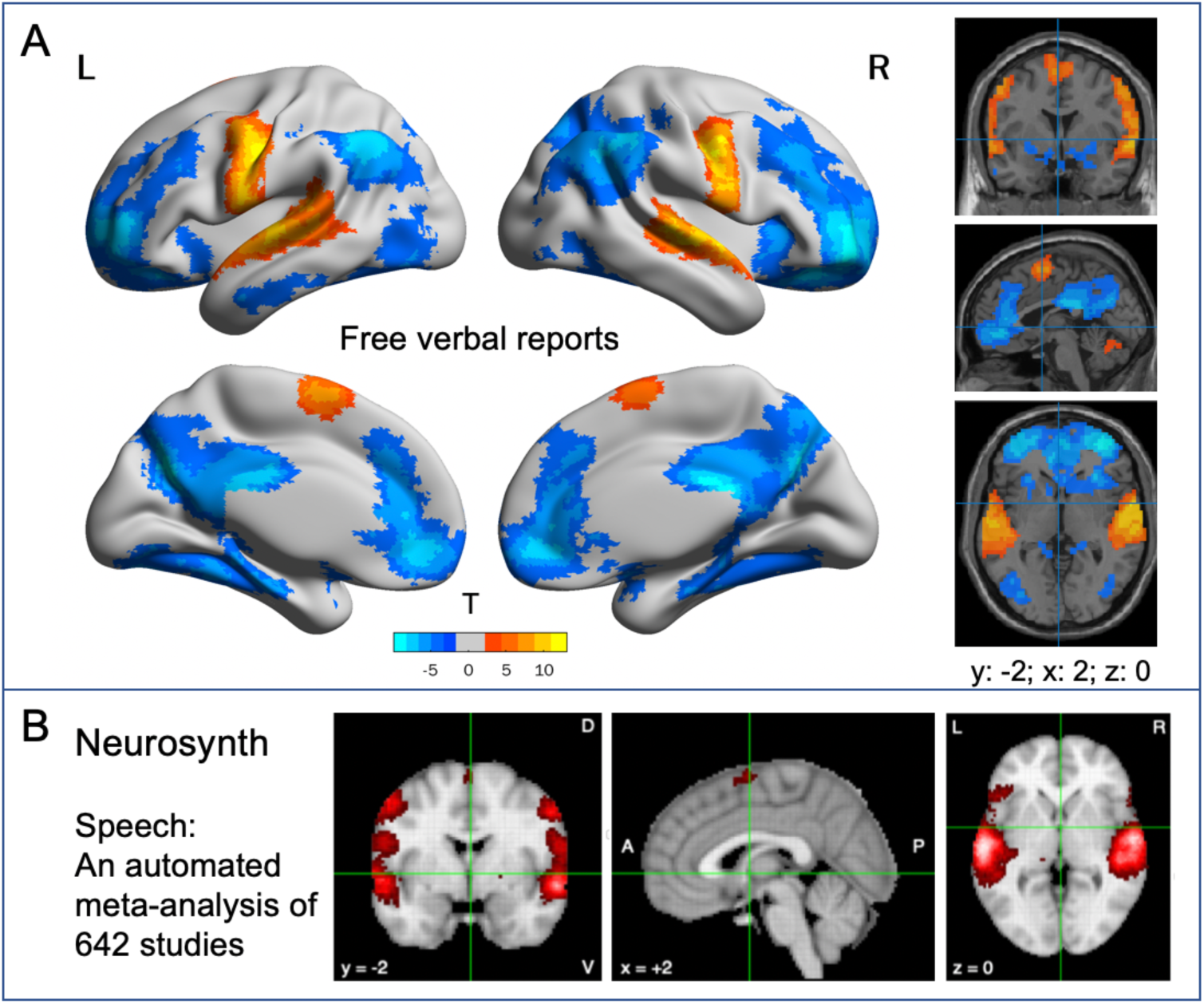
Brain activation patterns during free verbal reports. A) The pattern of activation in the free verbal report. B) Meta-analysis of 642 studies for the term “speech” based on Neurosynth.

### 3.3 The results of divergence in thought content

To test whether there was a relationship between brain activities related to self-generated thought reporting and thought content, we calculated correlations between the indicator of thought content divergence and brain activation maps. During the free verbal report stage, participants reported an average of 22 self-generated thought episodes. The frequency distribution can be found in Fig. 3C. We used natural language processing (NLP) to calculate the divergence of thought content over time (between two five-minute segments). We then calculated the correlation between the divergence indicator and the beta-value of the free verbal report stage at the voxel and Schaefer 400-parcels levels, respectively (Fig. 3). Both results showed that the divergence of thought content was significantly correlated with many brain regions (Fig. 3A, B): in addition to brain regions with significant positive activation in the reported activation results, we also detected a large number of brain regions with significant negative activation. Thus, the significantly deactivated brain areas in the report activation analysis reflect the content of thought.

**Fig. 3.**
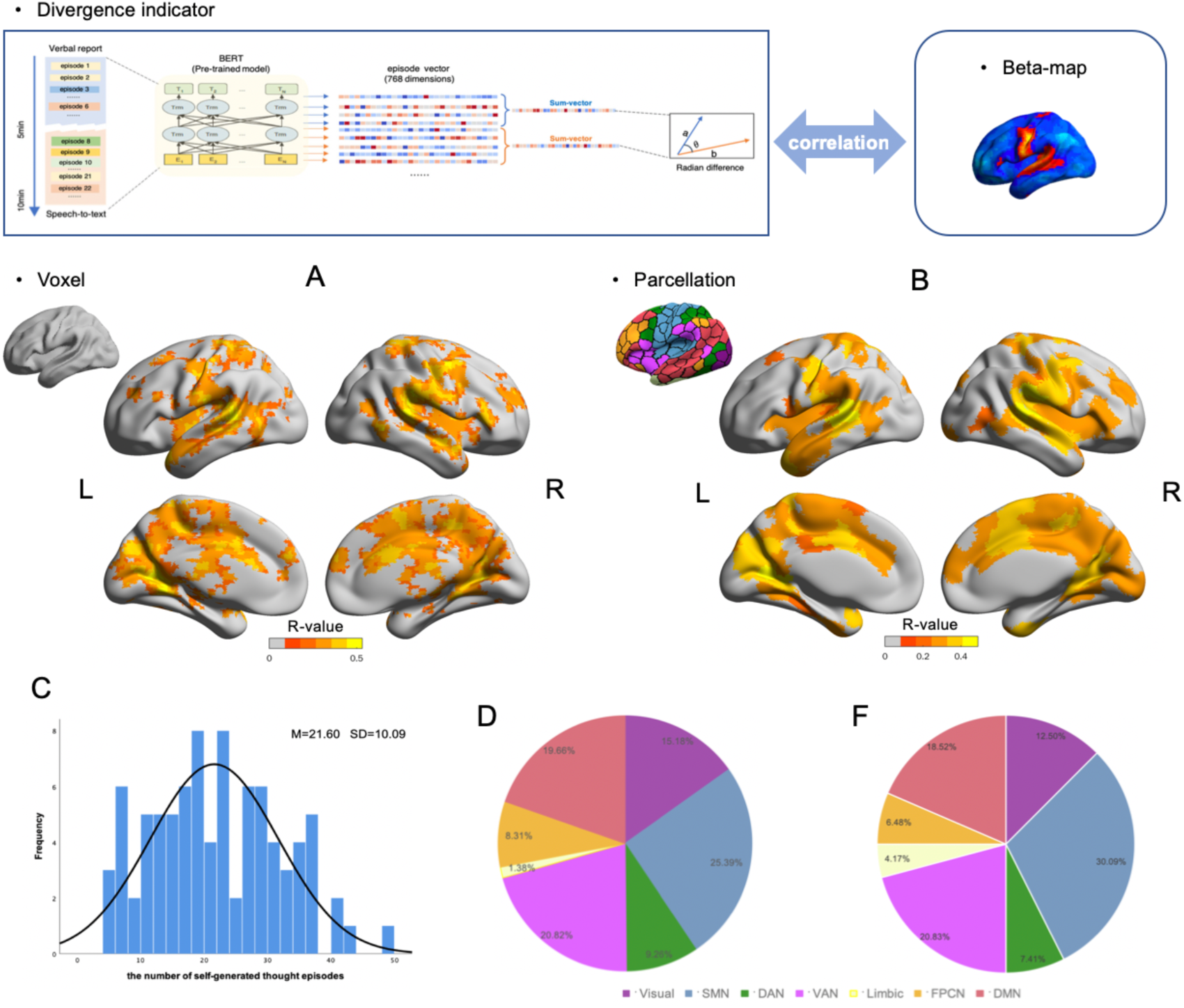
Results of brain representations of divergence in thought content. Correlation results between the divergence indicator in thought content and the beta-map in the free verbal report stage based on voxels (A) and Schaefer 400-parcels (B). Multiple comparisons correction: Threshold-Free Cluster Enhancement (TFCE) with permutation test (Voxel-wise); FDR correction, q <0.05 (Schaefer 400-parcels). The frequency distribution of self-generated thought episodes (C). Significant voxels (D) and parcels (F) were distributed across seven Yeo networks. The ratio of the number of significant voxels/parcels divided by the total number of significant voxels/parcels.

We examined these results by reference to the corresponding Yeo networks and calculated the ratio of the number of significant voxels/parcels divided by the total number of significant voxels/parcels (Fig. 3D, F). At the voxel level, the largest proportion of significance was found in the somatomotor network (SMN, 25.39%), followed by the ventral attention network (VAN, 20.82%), the DMN (19.66%), the visual network (Visual, 15.18%), dorsal attention network (DAN, 9.26%), frontoparietal control network (FPCN, 8.31%) and limbic network (Limbic, 1.38%). The parcel results were similar to those at the voxel level.

### 3.4 Representational similarity analysis results

We detected direct neural representations of self-generated thoughts in the “resting-state” scan through RSA at three different scales: a voxel-wise analysis with a whole-brain searchlight method; a region-level analysis with the Schaefer 400-parcels; at the systems level with the seven Yeo networks and language subsystems (Fig. 1).

#### 3.4.1 Voxel-wise whole-brain level RSA results

Whole-brain searchlight analysis was performed, in which neural RDMs were computed in a specific cubic centered on each voxel of the brain, and their relationships to behavioral RDMs were investigated. Given the difference in the word count of each reported self-generated thought episode, all subsequent RSAs used partial Spearman correlation, i.e., with the sentence length RDM as a covariate. The RSA searchlight mapping for the thought RDM yielded many significant regions that were widely distributed in both hemispheres (Fig. 4A) and involved all seven Yeo networks (Fig. 4B), with sentence length RDM covaried. The largest proportion of activity was in the DMN (27.20%), followed by the FPCN (16.21%), Visual network (14.37%), VAN (13.03%), SMN (12.65%), DAN (10.61%) and Limbic network (5.93%) (Fig. 4C).

**Fig. 4.**
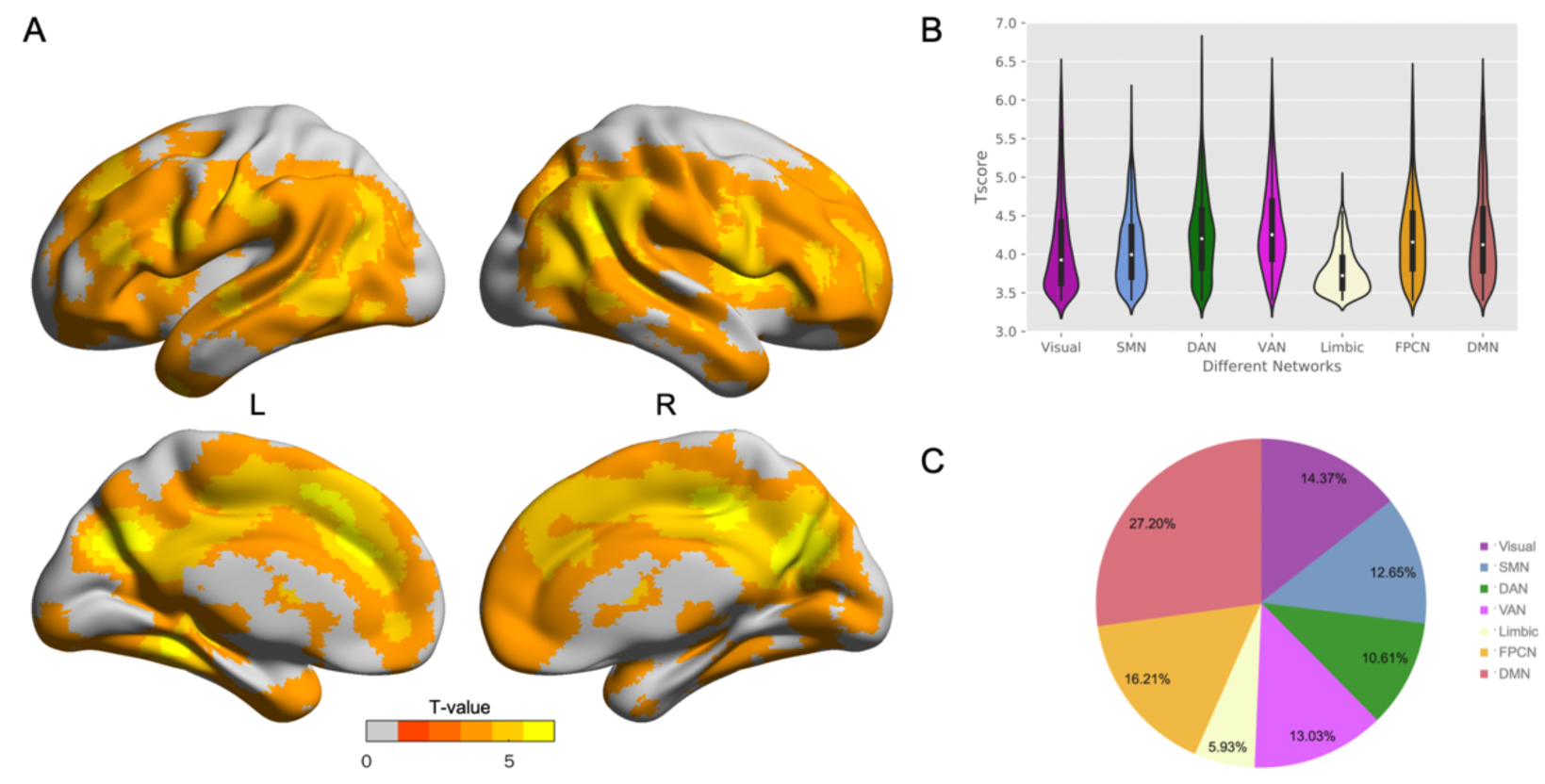
RSA results of reported self-generated thought content. A) Voxel-wise searchlight showed regions whose activity patterns had a significantly positive correlation with the reported thoughts. Partial Spearman correlation was calculated by controlling for sentence length. Multiple comparison corrections: GRF (voxel-level p < 0.001, cluster-level p < 0.05). B) Distribution of significant voxels in the seven Yeo networks. C) Ratio of the number of significant voxels in each network. Visual: visual network; DAN: dorsal attention network; Limbic: limbic network; DMN: default mode network; SMN: somatomotor network; VAN: ventral attention network; FPCN: frontoparietal control network.

#### 3.4.2 Region-level RSA results

We performed whole-brain RSA analysis based on the 400-parcels functional atlas of Schaefer et al. controlling for sentence length RDM in Spearman’s partial correlation. Two hundred and four parcels survived Bonferroni correction (p = 0.05/400), distributed throughout the Yeo networks (Fig. 5): 33.82% in the DMN, 16.67% in the VAN, 14.71% in the FPCN, 14.71% in the SMN, 8.82% in the DAN, 7.84% in the Visual network and 3.43% in the Limbic network.

**Fig. 5.**
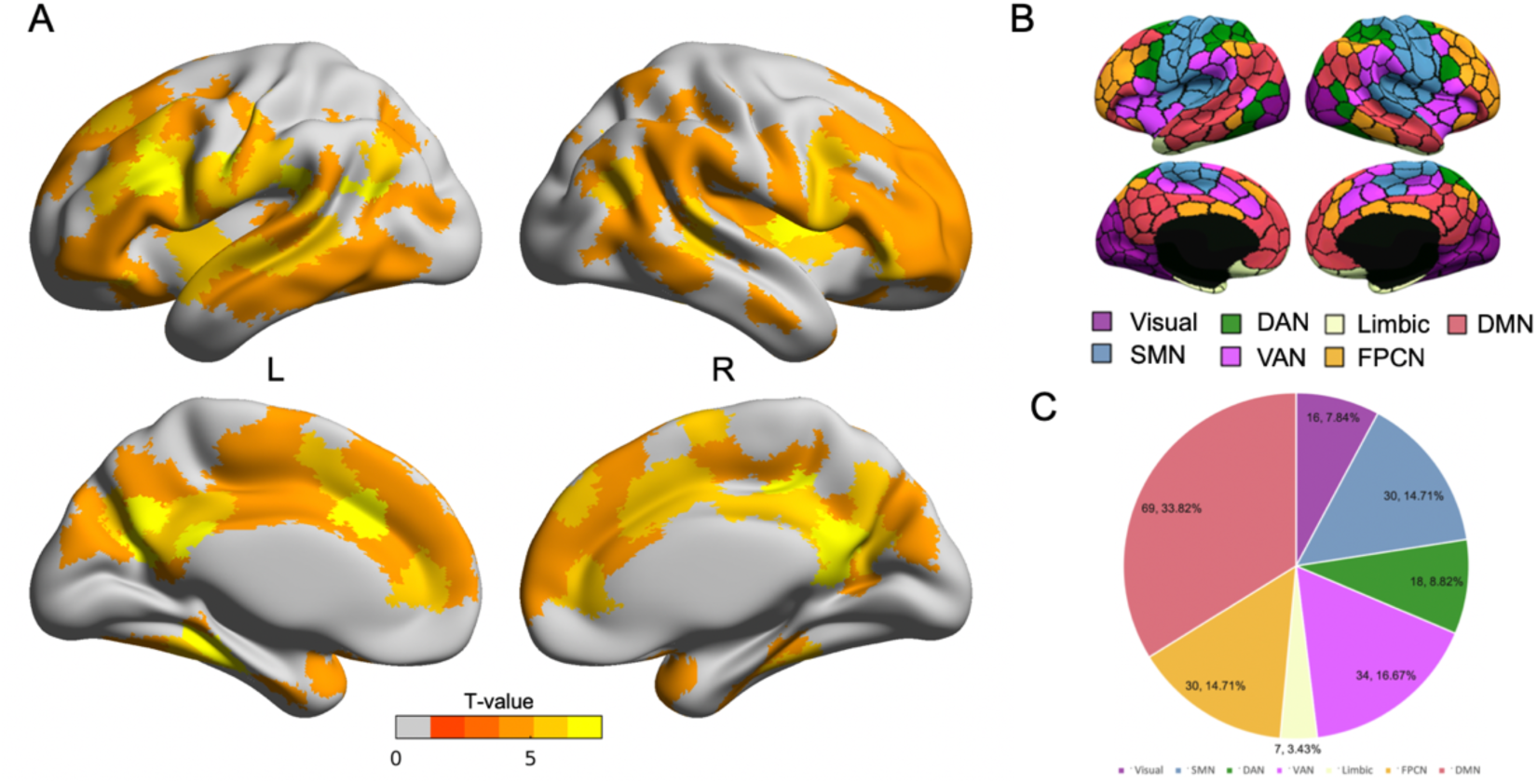
RSA results based on the 400-parcel functional atlas of Schaefer et al. A) Region-wise RSA showed parcels whose activity patterns had a significantly positive correlation with the reported thought. The partial Spearmen correlation was calculated by controlling for sentence length. Bonferroni correction was used for multiple comparisons (0.05/400). The results were shown at p < 0.05/400. B) 400-parcels were matched with the Yeo seven networks. C) The proportion of significant parcels that overlapped with the Yeo networks.

#### 3.4.3 System-level RSA results

The Yeo network masks were used to perform RSA and all seven networks’ neural RDM significantly correlated with the semantic feature RDM. The group statistic results are shown in Table 1. All p-values survived Bonferroni correction (0.0071).

**Table 1.** RSA results based on the Yeo networks

| Yeo seven-networks | t | p |
| --- | --- | --- |
| Visual network | 3.876 | 0.0002087 |
| Somatomotor network | 4.989 | 0.0000032 |
| Dorsal attention network | 4.132 | 0.0000840 |
| Ventral attention network | 5.654 | 0.0000002 |
| Limbic network | 3.549 | 0.0006327 |
| Frontoparietal control network | 4.505 | 0.0000210 |
| Default mode network | 5.622 | 0.0000002 |

Note: Sentence length RDM was a covariate. Bonferroni correction $p < 0.05/7$ .

### 3.5 Validation analysis

Validation analysis was performed to rule out possible confounders. First, head motion during the verbal report stage was below the criterion (mean FD_Jenkinson < 0.2 mm) for all analyzed participants. We correlated head motion and verbal report stage beta-values. No brain regions were significantly correlated with head motion. Second, we randomly permuted the order of participants’ reported thought episodes to construct the semantic RDM in random order. Then this RDM was used to perform voxel-wise-level searchlight RSA, region-level RSA, and system-level RSA (Table S4), respectively. We did not find any significant results in any of the levels of analysis.

Third, the RSA results using the beta-map for each episode were in general agreement with the currently used mean-BOLD maps. The RSA searchlight mapping for the thought RDM also yielded many significant regions and involved all seven Yeo networks (Fig. S4). The largest proportion of activity was in the DMN (24.96%), followed by the FPCN (20.38%), Visual network (11.83%), VAN (11.77%), SMN (14.38%), DAN (10.42%) and Limbic network (6.25%) (Fig. S4). The Region-level RSA results were also distributed throughout the Yeo seven networks (Fig. S5, Fig. S6). In addition, the RSA results based on the Yeo seven networks survived Bonferroni correction (Table S3). More details can be found in the supplementary material (2. The RSA based on beta-maps).

Fourth, we also calculated another indicator reflecting fluctuations in thought content and calculated the relationship between this indicator and brain activity. The details of the calculation are described in the Methods section. The results were consistent with the results of the divergence indicator; both voxel-level and parcel-level analyses showed that fluctuations in thought content were significantly correlated with many brain regions, involving all Yeo seven networks. The detailed results are shown in Figure S10.

## 4. Discussion

Investigators have suggested that considering ongoing experiences during the resting-state is important for the development of resting-state fMRI, however, few studies have investigated self-generated thoughts directly. To our knowledge, this is the first study to combine online experience sampling with dynamic neural activity measures. Here, we used the think-aloud fMRI method to collect participants’ real-time self-generated thoughts during “resting-state” scans. Our primary research question was to explore the direct neural representation of ongoing experiences in the resting-state. In addition, we also explored the feasibility of the think-aloud fMRI approach for studying ongoing experiences, including elucidating brain activation during verbal reports and whether thought content was significantly related to brain activity. Finally, we used RSA to explore direct brain representations of ongoing experiences. Specially, we used NLP to quantify each self-generated thought episode into a 768-dimension fixed-length vector to construct semantic feature RDM. To clarify how self-generated thoughts manifest in the brain, we investigated the neural correlations between reported thoughts and their corresponding brain activity by RSA at three scales. Our main finding was that “resting-state” self-generated thoughts involved a wide range of brain regions across all seven Yeo networks.

One of the factors that may account for the modest reliability of resting-state fMRI is the inherent complexity of ongoing mental activity during scanning (Morcom and Fletcher, 2007). The classical resting-state fMRI design is simple (“look at the ‘+’ fixation point and stay awake” or “close your eyes and rest”), making it difficult to influence and determine participants’ mental states during scans (Power et al., 2014). Participant self-generated thoughts may include planning the future, recalling the past, reflecting, and ruminating. Clearly, the so-called “resting-state” is not a complete standstill of the brain (Buckner, 2012), but rather reflects a mixture of mental activity and intrinsic functional organization (Finn, 2021). Likely because we pay little attention to observing these ongoing mental activities during resting-state scans, the interpretability of resting-state fMRI is inferior (Gonzalez-Castillo et al., 2021). Resting-state results cannot be simply attributed to the brain’s intrinsic activity. Thus, examining ongoing experiences during resting-state scans is increasingly important. However, such research has largely relied on retrospective introspection in which participants were asked to assess the characteristics of self-generated thoughts along certain dimensions throughout the resting-state scan. This method provides a window into ongoing experiences in the resting-state but because of methodological limitations, we are still unable to definitively study self-generated thoughts during resting-state scans. In this study, we used an immediate think-aloud fMRI approach to collect real-time ongoing experiences during “resting-state” scanning. Moreover, we combined NLP to directly quantitatively analyze self-generated thoughts, avoiding the lag of retrospective methods and reducing introspection, as well as expanding the study of the content features of self-generated thoughts (open to exploration without being limited to predefined thought dimensional characteristics).

However, a major consideration in applying the think-aloud method to MRI relates to the potential of speaking to increase head motion, and to the impact of speaking on brain activity. We took some measures to control participants’ head motion (see materials and methods). We found that most participants were able to control themselves well, with only 3 out of 101 participants having head motion that exceeded criterion (mean FD_Jenkinson < 0.2 mm) (Fig. S2). In addition, we calculated correlations between our head motion index (mean FD_Jenkinson) and individual brain activations (beta-maps) and did not find any brain regions significantly associated with individual head motion. As to whether speaking has a significant effect on individual brain activity, we expected individuals would significantly activate brain areas associated with speech when they reported their thoughts (Fig. 2). Furthermore, we designed a control condition, i.e., a cue report (Fig. S7) to test language processing. The control verbal report stage was a block design, in which when participants saw the “report” clue word, they needed to report any thoughts or images that occurred in their minds during the next 20 seconds, and when they saw the “don’t report” clue word, they did not need to report their thoughts during the next 20 seconds (Fig. S7). If there were no thoughts/images in the participant’s mind at the stage where reporting was required, then they did not need to report. Participants were required to report only when thoughts/images were present during the reporting phase. More information about the control conditions is in the supplementary material. In this block design, the non-reporting phase is the classical resting-state and the reporting phase is the immediate think-aloud phase. The order of reporting and non-reporting blocks was randomized. We assumed that individuals would produce a series of thoughts and mental images during both the non-reporting and reporting blocks with the only difference between the blocks being whether they spoke or not. By comparing the free reporting and non-reporting blocks we sought to examine the impact of speaking. In comparing the reporting phase with the non-reporting phase, we found this brain pattern is largely consistent with that of the free verbal report stage (Fig. 2, Fig. S8). Furthermore, we obtained the brain regions required for “speech” based on Neurosynth (https://www.neurosynth.org) (Fig. 2) and found that the brain regions that were significantly positively activated in the activation pattern maps of the free speech and control stages were included in the brain regions required for “speech”. Verbal reports do significantly activate the brain areas associated with language processing. Notably, consistent with the free verbal report stage, the language areas were significantly activated, while many brain regions were deactivated. Brain regions deactivated during the free verbal report stage involved the default mode network, which is usually deactivated during tasks with a large cognitive load (Buckner et al., 2008). Here speaking out thoughts (think-aloud) may involve more cognitive load than silent thinking (without speaking), that is, the cognitive load may be significantly greater in the reporting phase than in the non-reporting phase. However, more research is needed to understand this deactivation phenomenon.

Researchers have begun to emphasize the importance of the dynamic nature of thought as proposed by the Dynamic Framework (Andrews-Hanna et al., 2020; Christoff et al., 2016). Deliberate (cognitive control) and automatic constraint (sensory and affective salience) influence the contents of thought and its transition over time. Unconstrained thought/freely-moving thought has relatively low constraints (O’Neill et al., 2021). The frequency of participants’ unconstrained thoughts has been found to be influenced by the difficulty of completing a task (Brosowsky et al., 2021). Specifically, participants engaged in more unconstrained thoughts when completing easy tasks than when completing difficult ones. Researchers (O’Neill et al., 2021) have questioned whether unconstrained thought based on task context studies reflects the lack of goal guidance, in other words, is it consistent with the concept proposed by Christoff et al? What is the nature of freely moving thought? It is at rest, i.e., in the absence of an externally imposed task, that people are most likely to experience unconstrained thought and exhibit a natural flow of thought. We told participants to relax and rest as much as possible without deliberately engaging in thinking and to let their minds move as freely as possible. According to the dynamic framework, thoughts in the resting-state arise and proceed in a relatively free, unconstrained fashion. We followed the nature of the flow of thought over time and processed the thoughts reported by participants according to topic similarities. We calculated metrics to characterize the variability of thought content. The divergence of thought content was the index of variation of thought content across two time periods calculated from the processed texts. This metric calculates the overall difference in thought contents across the two five-minute periods. A participant may switch (freely moving) her thoughts many times, or he may also switch among a few topics repeatedly. In contrast, a participant may switch his thought only a few times, but each time onto a separate topic, so the variability of the calculated overall thinking content may be large. We believe that the divergence indicator can, to some extent, reflect the breadth of topics to which individuals constrained the content of their thoughts. People with high rumination may tend to automatically limit their thoughts to fewer topics or events (Christoff et al., 2016; Treynor et al., 2003). In the other words, they generally stick to similar thoughts (DuPre and Spreng, 2018). Our previous study has demonstrated that this indicator was significantly negatively correlated with rumination, suggesting that this index of thought content divergence reflects the extent to which thought content was constrained over time (Li et al., 2021). To further illustrate the brain regions associated with the content of thought, we calculated correlations between indicators reflecting differences in thought content and brain activity (Fig. 3). Specifically, we calculated the correlation between the divergence indicator and the beta-value of the free verbal report stage at the voxel and Schaefer 400-parcels levels, respectively (Fig. 3). The results show that a wide range of brain regions are associated with the content of thought. In addition, we also calculated another indicator of thought content, called the fluctuation indicator, by first calculating the difference between each thought episode within a 10- minute phase and the average vector for that phase, after which we obtained the coefficient of variation (Fig. S10). The fluctuation indicator also reflects the fluctuation of thought content over time and contains information about participants’ constraints on thought content, but unlike the divergence indicator, it reflects each switch (moving) in content and indicates the magnitude of the oscillation of the thought content switch over time. Correlation analysis with brain activity also showed that thought content was significantly correlated with a wide range of brain regions (Fig. S10). The correlation analysis between these two metrics and brain activity identified, to some extent, the brain regions associated with thought content.

In particular, we computed neural response patterns of self-generated thoughts in the think-aloud fMRI using RSA on three neural scales and found that the representation of self-generated thoughts in the “resting-state” is diffuse: it is not just the DMN that characterizes self-generated thoughts but a wide range of regions throughout the brain are involved, including all seven Yeo networks. This result confirms previous findings. A meta-analysis by Fox et al. found that while spontaneous thought processes were significantly correlated with the DMN, many non-DMN regions also had consistent activity such as secondary somatosensory cortices, lingual gyrus, and insula. Thus, they emphasized that the DMN was not sufficient to adequately capture the neural basis of self-generated cognitive process and that regions of non-DMN networks, such as FPCN, were as important to spontaneous thoughts as the DMN (Fox et al., 2015). Moreover, functional connectivity (FC) among networks plays a role in the characteristics of self-generated thoughts at rest. Wang et al. calculated the correlation between self-generated thoughts during resting-state scans and large-scale network organization based on the retrospective self-report method (Wang et al., 2018a). The results showed that the four different dimensions of self-generated thought at rest correlated differently with different FCs. For example, experiences focused on “personal importance” were associated with reduced FC within the attention and control system, while “purposeful” experiences were correlated with lower FC between DMN and limbic network. Furthermore, given the increasing importance of the three subsystems of the DMN in the field of self-generated thought (Andrews-Hanna et al., 2014; Buckner et al., 2008), we also conducted RSAs based on the three subsystems. We defined a total of 24 regions of anatomical interest based on 17 network parcellations derived by Yeo from the data of 1000 young healthy individuals (Yeo et al., 2011), and further divided into 3 DMN subsystems (Fig. S11) (Andrews-Hanna et al., 2014; Dixon et al., 2017). The results showed that all three subsystems were significantly correlated with the reported thoughts (core subsystem: t = 5.563, p < .0001, the dorsal medial prefrontal cortex (DMPFC) subsystem: t = 5.072, p < .0001, the medial temporal lobe (MTL) subsystem: t = 4.443, p < .0001). The three DMN subsystems promote different self-generation thoughts and have close interaction (Andrews-Hanna et al., 2014; Buckner and DiNicola, 2019; Chen et al., 2020). Therefore, the complexity of the content of resting state ongoing experiences was also reflected in the interaction between the three subsystems of the default network.

Following the dynamic nature of the mind and the brain itself, based on the RSA approach, we obtained a modal of the spatial representation of the brain inherent to the ongoing experience of the resting state. As expected, the mental activity of the resting state involves a wide range of brain regions. Moreover, this may reflect the spatial features of the “common currency” proposed by Spatiotemporal Neuroscience (Northoff et al., 2020). Self-generated thought at “rest” is complex and variable (Wang et al., 2018b), its form includes images, words, etc., and its content includes temporal dimensions, social dimensions, emotional valence, etc. (Gorgolewski et al., 2014). In this study, in addition to the DMN and FPCN, we found a substantial proportion of significant voxels in the visual network, which is consistent with individuals reporting that much of inner experience in the task-free state involves visual imagery (Delamillieure et al., 2010). Relatedly, retrospective assessment after the free verbal report stage scan showed that participants related many visual imagery experiences (Fig. S1, Table S2). Visual networks have an important role in visual imagery (Ganis et al., 2004), and researchers have demonstrated that visual areas are associated with the emergence of visual, non-verbal features of self-generated thoughts; moreover, visual thoughts were associated with more positive valence (Raij and Riekki, 2017). Except for visual thoughts, a large portion of self-generated thoughts take the form of internal verbal behavior (thinking in words without overt vocal production) (Delamillieure et al., 2010; Raij and Riekki, 2017). There were three modules based on resting-state that were related to semantic processing (Fig. S9A), located in the default mode network, the left perisylvian network, and the left frontoparietal network (Xu et al., 2016). The significant voxels from the whole-level searchlight RSA yielded proportions of 47.16% for the DMN Module, 36.28% for the FPN Module, and 16.56% for the PSN Module (Fig. S9C). Meanwhile, the RSA results based on three modules showed significant correlations between the three networks and the reported thoughts (Fig. S9A). The activity of attention networks including DAN and VAN may reflect the transition between internally driven and externally driven resting-state experience. Although there was no explicit external task during the resting-state scan, scanner noise was still present, and participants still needed to keep their eyes open to gaze at the fixation point. Self-generated thoughts focused on “personal importance” were associated with the attention network (Wang et al., 2018a). The SMN may represent an individual’s processing of external and internal sensations, including scanner noise and self-body somatomotor experiences. Moreover, the somatomotor network was important for language processing (Pulvermüller and Fadiga, 2010). Finally, the limbic network embodies emotional experience in self-generated thoughts at “rest”. Self-generated thoughts involve emotional information and are linked to mood state (Andrews-Hanna et al., 2013; Killingsworth and Gilbert, 2010; Ruby et al., 2013), such that healthy individuals think about more positive things while depressed individuals think about more negative events. Furthermore, the FC of the limbic network to other networks was found to be related to different forms of experiences at rest. For example, emotional experiences were associated with higher FC between limbic and VAN; visual experiences were associated with stronger connectivity between visual and limbic networks (Wang et al., 2018a).

We note a few points about conceptual aspects. First, research in the field of mind wandering that focuses on task-independent cognitive processes may provide a pathway for studying self-generated thoughts in the “resting-state”. Indeed, verbalizing the content of one’s own thoughts (think-aloud) is not precisely equivalent to mind wandering, whether it be freely mind wandering or task-unrelated mind wandering. Think-aloud is more closely related to freely mind wandering, but with a higher cognitive load to the requirement to speak out loud. Our previous study (Li et al., 2021), together with others (Raffaelli et al., 2021) showed the characteristics of self-generated thoughts during think-aloud are similar to those during the resting-state. Therefore, the results of the present study can have implications for the field of self-generated thoughts but with caution.

Second, researchers have emphasized that self-generated thoughts can occur intentionally (deliberate) or unintentionally (spontaneous) and both kinds of mind wandering were caused by different processes (Seli et al., 2015; Smallwood and Schooler, 2015). Research on whether self-generated thought was intentional was usually based on the task context, where thought probes (self-reports) were used to confirm whether participants’ mind wandering was intentional or unintentional. Researchers have demonstrated that both forms of mind wandering can exist simultaneously and that unintentional mind-wandering is more frequent than intentional mind-wandering, regardless of the type of task being performed (Seli et al., 2016a; Seli et al., 2016b). In addition, these two forms of thought are differentially related to task difficulty; intentional mind wandering rate was higher in an easy task than in a difficult task, whereas unintentional mind wandering rate was lower in an easy task than in a difficult task (Martínez-Pérez et al., 2021; Seli et al., 2016a). We did not measure whether each self-generated thought episode was generated intentionally or unintentionally. Based on the findings described above, we believe that self-generated thoughts in the resting-state contain thoughts generated in both ways and that more thoughts were generated intentionally and fewer were generated unintentionally compared to the task condition. In addition, the boredom of the resting-state and the fact that we asked participants to report their thoughts may have facilitated the generation of intentional thoughts.

Third, cognitive control was closely related to self-generated thoughts. People with good cognitive control produce more mind wandering when the environment is undemanding (Kane et al., 2007; Levinson et al., 2012; Smallwood and Schooler, 2015). In our study, the cognitive load of think-aloud was much lower compared with most task paradigms. This likely enhanced the frequency of self-generated thoughts by our participants. More research is needed to explore differences between self-generated thoughts in the resting-state and task states.

The main contributions of this study are that we: 1) investigated the think-aloud fMRI paradigm (i.e., combining the think-aloud method with neuroimaging) to identify the neural mechanisms of the stream of thought during “resting-state”; 2) analyzed data by combining the NLP method with the searchlight neuroimaging technique; 3) proposed and provided evidence that self-generated thoughts involve all brain networks, rather than being restricted to a single or a few networks. By directly combining the spontaneity and dynamic of thought with the spontaneity and dynamic of brain activity, we empirically obtain an intrinsic brain spatial pattern of spontaneous activity in the brain’s resting state. This study highlights the importance of attention to ongoing experiences in the resting state, clarifies the internal spatial model of brain spontaneous activity, and provides experimental support for Spatiotemporal Neuroscience (Northoff et al., 2020). In addition, the brain atlas of intrinsic activity in the resting state obtained in the present study can clarify the spatial structural features of brain spontaneous activity and promote the development of brain spontaneous activity research. Methodologically, this study demonstrated the potential of integrating the think-aloud method, natural language processing, and representational similarity analysis with fMRI for studying the connection between stream of thought and spontaneous brain activity in the resting state.

## Limitation and future directions

The current study used the think-aloud method to directly quantify the form of brain representations of self-generated thoughts during approximate “resting-state” fMRI. However, the current study is a preliminary exploration, with some limitations. First, the use of think-aloud to study self-generated thoughts is an initial means of revealing the multifaceted features of thoughts while allowing for quantitative analyses. However, the “resting-state” scan that is performed during think-aloud is not precisely equivalent to a simple resting-state scan. Therefore, more natural methods should be devised that can better expand the study of resting-state self-generated thoughts. Second, our study empirically demonstrated that self-generated thoughts involve a wide range of brain regions and are not limited to a single or a few networks. However, it is unclear how the different content features of self-generated thoughts relate specifically to the corresponding networks. Addressing this aspect will require clarifying the specific characteristics of each reported thought, such as whether it belongs to past event recall or body sensation, etc. Examining reported self-generated thought episodes with labels will help us to clarify the specific brain activity patterns corresponding to different experience types. We recruited psychologically knowledgeable raters to rate the content dimensions of participants’ self-generated thought episodes, such as sentiment and social. Future work should further clarify the neural representations of the different contents of self-generated thoughts. Third, resting-state ongoing experience is complex; it is a mixture of multiple forms and contents of thought, including fully internally oriented thought as well as thought elicited by external stimuli, unconstrained thought as well as actively constrained thought, autobiographical memory or planning, interoceptive, different forms of representations of thought such as verbal, visual, and auditory, and different forms of thought such as rumination and reflection. The current research did not break down specific forms of thinking and various content features but rather encompassed them uniformly within the resting-state thought stream. More research is needed to explore the different forms and contents of resting-state thinking in more detail. Fourth, we did not collect physiological data such as participants’ respiratory and heart rates, therefore we were unable to model physiological fluctuations that could be bidirectionally modulated by neural activity. Fifth, we did not examine how the neural representation of self-generated thoughts might change across mental disorders. Obtaining real-time self-generated thoughts data with think-aloud fMRI across different psychiatric disorders is needed to help us to clarify how to control for the impact of ongoing experiences when comparing between groups, and to better interpret resting-state fMRI results.

## Conclusion

In conclusion, we directly and quantitatively demonstrated that self-generated thoughts during think-aloud fMRI were closely associated with a wide range of brain regions. This study highlights the need to account for the characteristics of self-generated thoughts when using resting-state fMRI. In particular, differences in resting-state fMRI self-generated thoughts should be considered when comparing groups, especially for groups with psychiatric disorders.

## Declaration of competing interest

The authors declare no conflicts of interest.

## Credit authorship contribution statement

**Hui-Xian Li**: Conceptualization, Methodology, Software, Formal analysis, Investigation, Data Curation, Writing - Original Draft, Writing - Review & Editing, Visualization, Funding acquisition. **Bin Lu**: Methodology, Investigation. **Yu-Wei Wang**: Investigation. **Xue-Ying Li**: Investigation. **Chao-Gan Yan**: Conceptualization, Methodology, Software, Writing - Original Draft, Writing - Review & Editing, Supervision, Project administration, Funding acquisition.

## Acknowledgment

We thank Prof. Francisco Xavier Castellanos for his comments and edits on our work. We appreciate Dr. Yanchao Bi for her comments and generously sharing their semantic masks. This work was supported by the Sci-Tech Innovation 2030 - Major Project of Brain Science and Brain-inspired Intelligence Technology (grant number: 2021ZD0200600), National Key R&D Program of China (grant number: 2017YFC1309902), the National Natural Science Foundation of China (grant numbers: 82122035, 81671774, 81630031), the 13th Five-year Informatization Plan of Chinese Academy of Sciences (grant number: XXH13505), the Key Research Program of the Chinese Academy of Sciences (grant NO. ZDBS-SSW-JSC006), Beijing Nova Program of Science and Technology (grant number: Z191100001119104), and the Scientific Foundation of Institute of Psychology, Chinese Academy of Sciences (grant number: E2CX4425YZ).

## Notes

### Competing Interest Statement

The authors have declared no competing interest.

### Summary of Updates

We added some theoretical notes and a description of the study's limitations in the revised manuscript. In addition, a new supplementary analysis has been added.

