## SupplementalMaterial for "Neural Representations of Self-Generated Thought during Think-aloud fMRI"

Contents:

1. Experimental Paradigms and behavioral analysis

2. The RSA based on beta-map

3. Control stage analysis

4. Other RSA results

List of Tables:

Table S1. Eight questions at the end of each stage

Table S2. Statistical results of paired t-tests on the content of self-generated thought from four dimensions.

Table S3. The beat-map RSA results based on the Yeo networks.

Table S4. RSA results for semantic RDM with random order based on the Yeo networks.

List of Figures:

Fig. S1. Comparisons of differences (two-tailed paired t-tests) across four dimensions of self-generated thought content.

Fig. S2. Distribution of head motion during free verbal report stage.

Fig. S3. Sentence length distribution based on all self-generated thought episodes.

Fig. S4. The searchlight RSA results of reported self-generated thought content based beta-map.

Fig. S5. Beta-map RSA results based on the Schaefer 400-parcels (FDR correction).

Fig. S6. Beta-map RSA results of thought RDMs based on the Schaefer 400-parcels (Bonferroni correction).

Fig. S7. Schematic diagram of the fMRI task.

Fig. S8. The activation results of the reporting vs. non-reporting condition in the control report stage.

Fig. S9. Results of three modules related to semantic processing of resting-state

Fig. S10. Results of brain representations of divergence in thought content.

Fig. S11. The three subsystems of DMN were used in the present study.

1. Experimental Paradigms related materials and behavioral analysis

**The retrospective rating questions.**

At the end of the scan participants rated their thought content (temporal, social, and emotional valence) and form (words and images) on a scale from 1 to 9 points.

Table S1. Eight questions at the end of each stage

| Dimension | Question |
| --- | --- |
| Temporal | My thoughts were about past. |
|  | My thoughts were about future. |
| Social | My thoughts were about myself. |
|  | My thoughts were about other people. |
| Emotion value | The content of my thoughts were positive events. |
|  | The content of my thoughts were negative events. |
| Mental experience form | My thoughts were in the form of images. |
|  | My thoughts were in the form of words. |

9-point Likert scale (1: not at all, 9: completely)

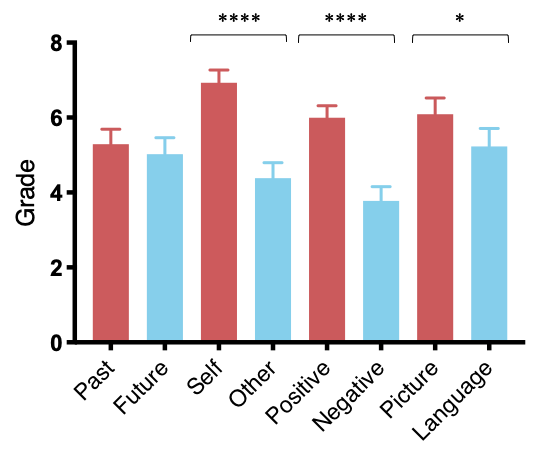

Fig. S1. Comparisons of differences (two-tailed paired t-tests) across four dimensions of self-generated thought content based on participants’ retrospective rating.

Table S2. Statistical results of paired t-tests on the content of self-generated thought from four dimensions (temporal: past vs. future; social: self vs. other; emotional valence: positive vs. negative; mental experience: picture vs. language).

| Dimension | t (df = 85) | p | Cohen’d | 95% CI |
| --- | --- | --- | --- | --- |
| Past-Future | 0.767 | 0.445 | 0.083 | -0.426, 0.961 |
| Self-Other | 7.789 | < .001 | 0.840 | 1.896, 3.197 |
| Positive-Negative | 6.693 | < .001 | 0.722 | 1.561, 2.881 |
| Picture-Language | 2.190 | 0.031 | 0.236 | 1.642, 0.236 |

**Participants’ head motion during the scanning**

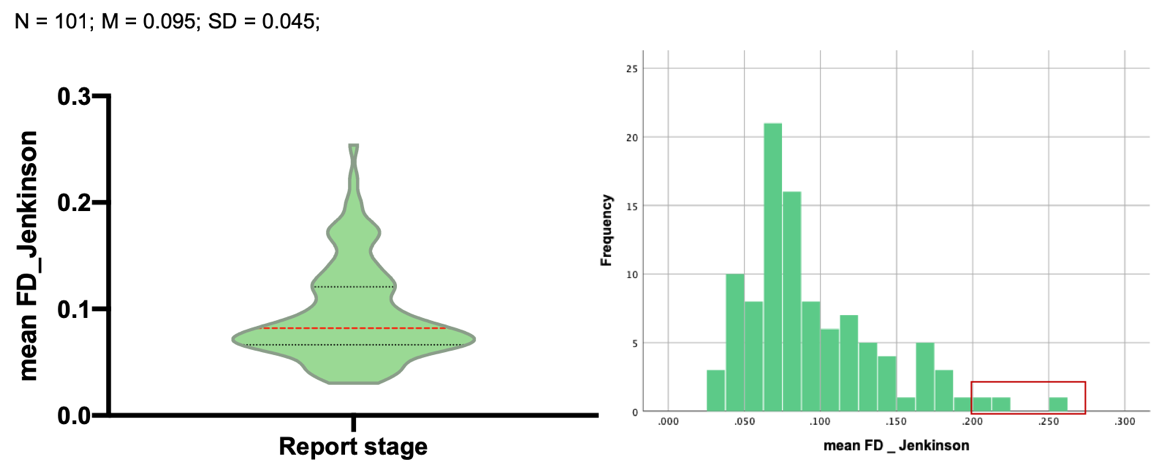

Fig. S2. Distribution of head motion during free verbal report stage. The sample size (N = 101) is the number of all participants who completed the scanning experiment. Three participants (red box) exceeded the head motion criterion (mean FD_Jenkinson > 0.2 mm).

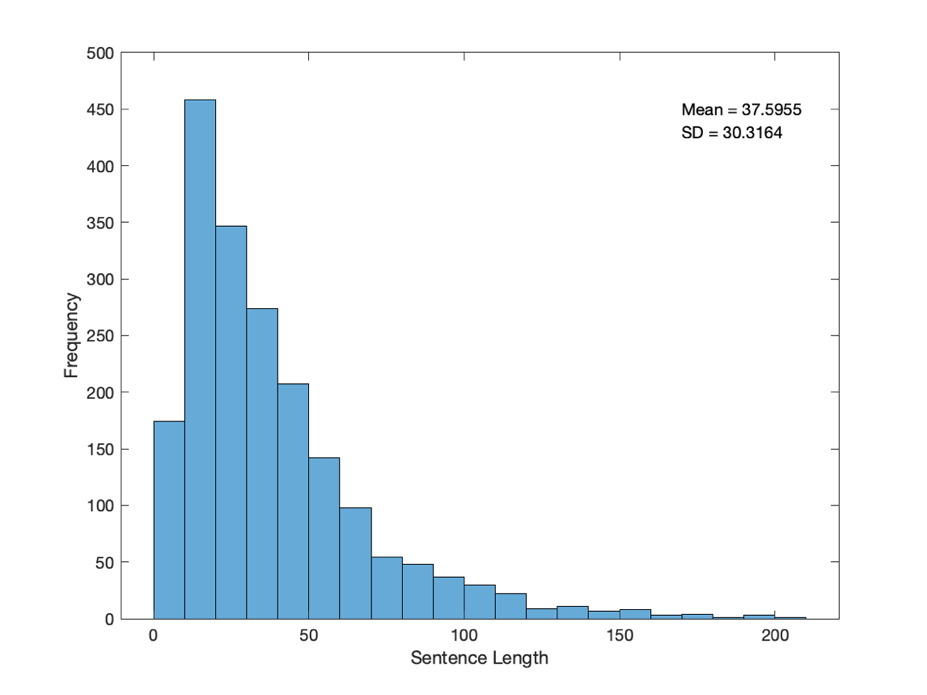

Fig. S3. Sentence length distribution was calculated based on all self-generated thought episodes.

2. The RSA based on beta-map

**Voxel-wise whole-brain level RSA results.**

The RSA searchlight mapping for the thought RDM yielded a large number of significant regions and involved all seven Yeo networks (Fig. S4), with sentence length RDM covaried. The largest proportion of activity was in the DMN (24.96%), followed by the FPCN (20.38%), Visual network (11.83%), VAN (11.77%), SMN (14.38%), DAN (10.42%) and Limbic network (6.25%) (Fig. S4).

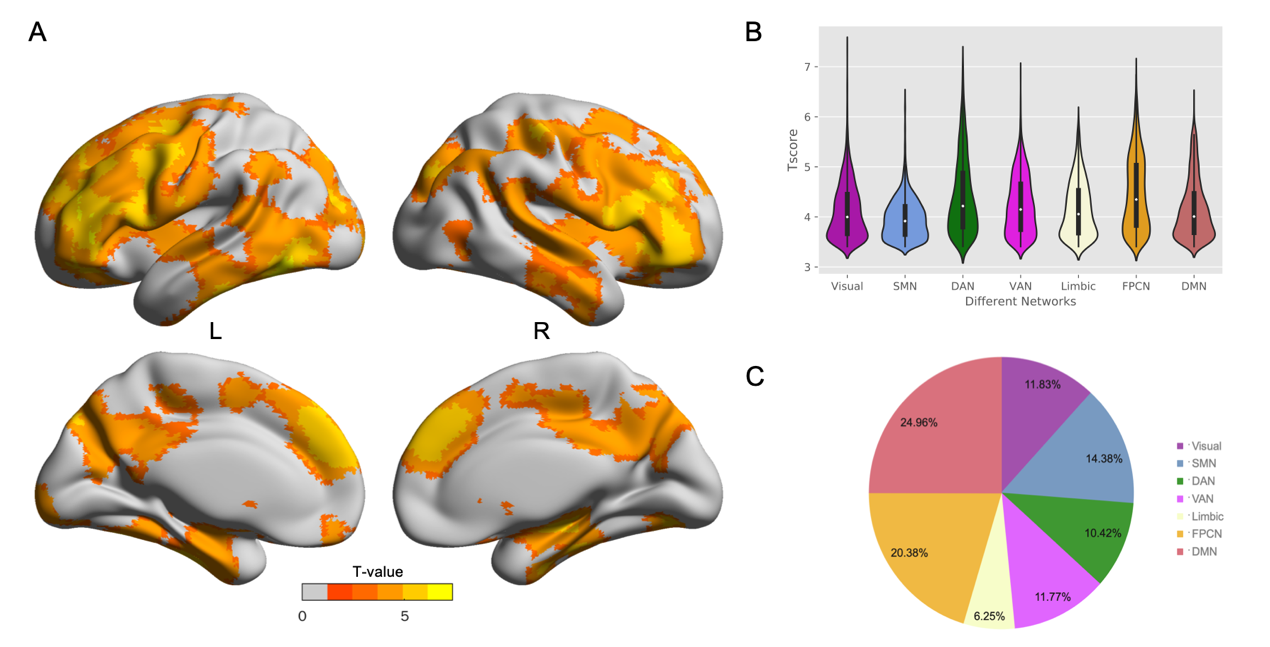

Fig. S4. The searchlight RSA results of reported self-generated thought content based beta-map. A) Voxel-wise searchlight showed regions whose activation patterns had a significantly positive correlation with the reported thoughts. Partial Spearman correlation was calculated by controlling for sentence length. Multiple comparison corrections: GRF (voxel-level p < 0.001, cluster-level p < 0.05). B) Distribution of significant voxels in the seven Yeo networks. C) Ratio of the number of significant voxels in each network. Visual: visual network; DAN: dorsal attention network; Limbic: limbic network; DMN: default mode network; SMN: somatomotor network; VAN: ventral attention network; FPCN: frontoparietal control network.

**Region-level RSA results.**

Two hundred and eighty-three parcels survived FDR correction (q = 0.05), distributed throughout the Yeo networks (Fig. S5): 24.38% in the DMN, 17.67% in the SMN, 15.90% in the FPCN, 13.43% in the Visual network, 12.01% in the VAN, 9.89% in the DAN and 6.71% in the Limbic network. When applying strict Bonferroni correction (p < 0.05/400), significant parcels were still distributed over all seven Yeo networks (Fig. S6).

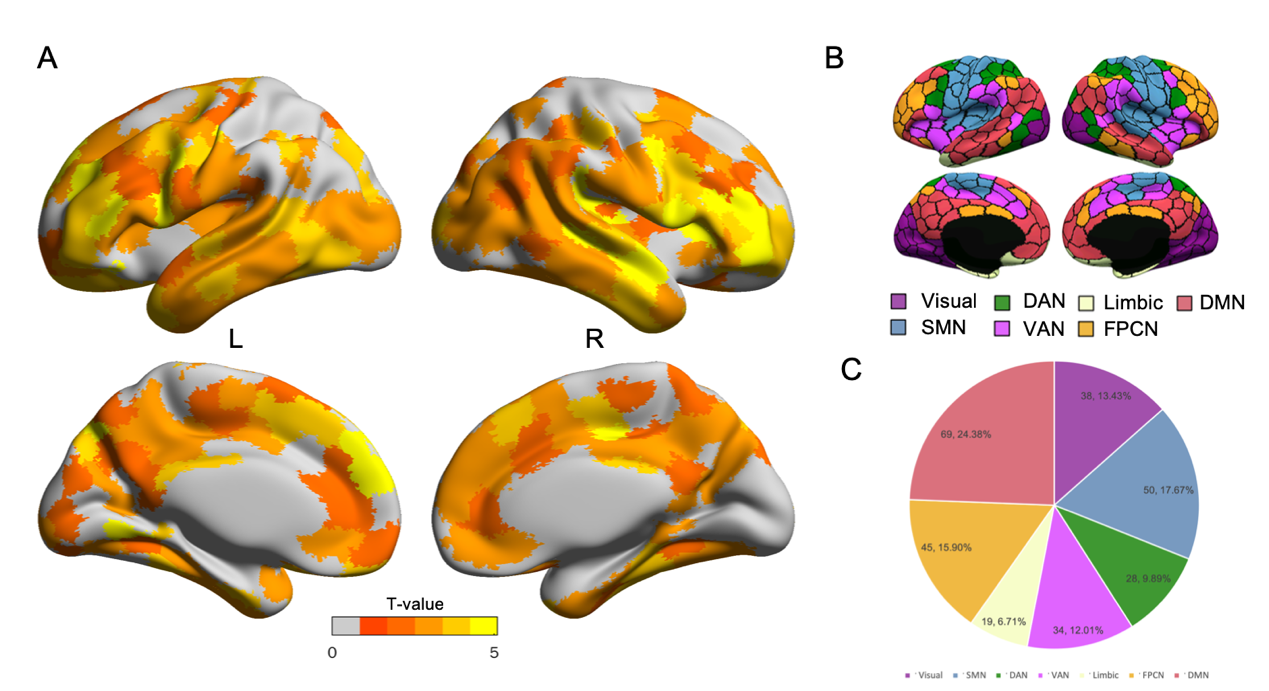

Fig. S5. Beta-map RSA results based on the 400-parcel functional atlas of Schaefer et al. A) Region-wise RSA showed parcels whose activity patterns had a significantly positive correlation with the reported thought. The partial Spearmen correlation was calculated by controlling for sentence length. FDR correction was used for multiple comparisons. Results were showed at the q < 0.05 level. B) 400-parcels were matched with the Yeo seven-networks. C) The proportion of significant parcels that overlapped with the Yeo networks.

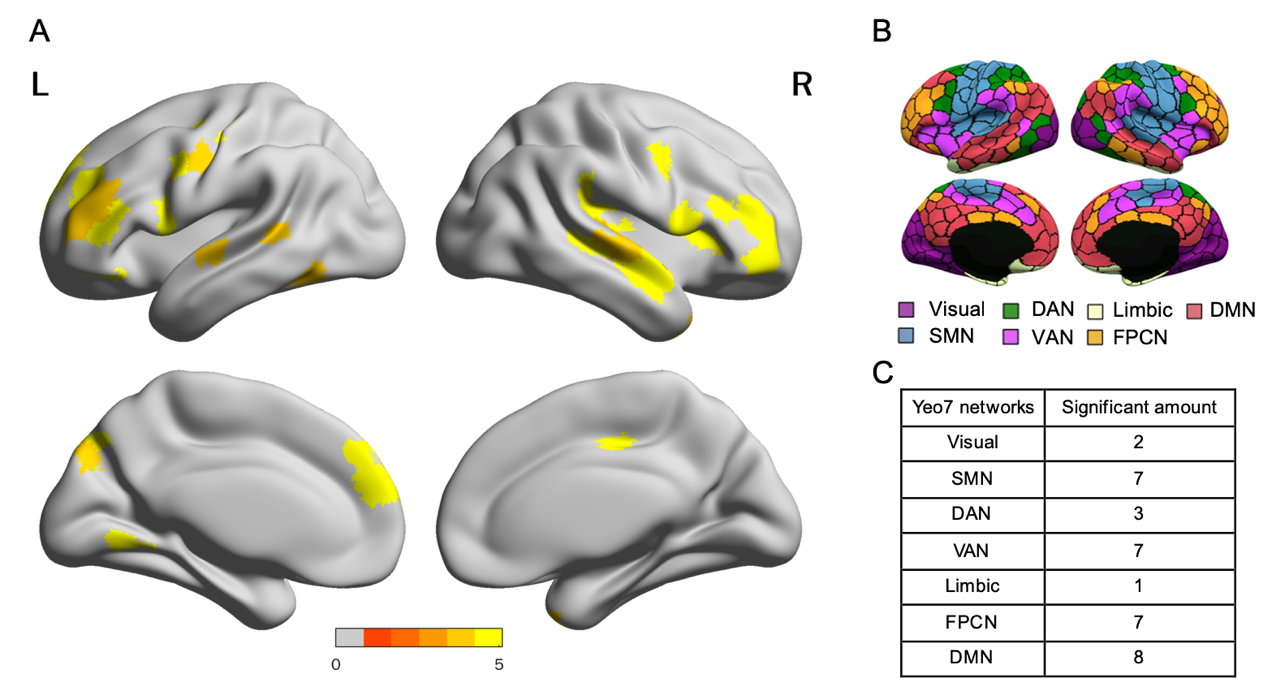

Fig. S6. Beta-map RSA results of thought RDMs based on Schaefer 400-parcels. We used Bonferroni correction. P < 0.05/400.

**System-level RSA results**

The seven Yeo network masks were used to perform RSA and all seven networks’ neural RDM significantly correlated with the semantic feature RDM (Table 1). All p-values survived Bonferroni correction (0.0071).

Table S3. The beat-map RSA results based on the Yeo networks

| Yeo seven-networks | t | p |
| --- | --- | --- |
| Visual network | 4.688 | 0.0000104 |
| Somatomotor network | 4.877 | 0.0000050 |
| Dorsal attention network | 3.915 | 0.0001818 |
| Ventral attention network | 5.257 | 0.0000011 |
| Limbic network | 2.838 | 0.0056813 |
| Frontoparietal control network | 5.554 | 0.0000003 |
| Default mode network | 5.327 | 0.0000008 |

Note: We used beat-map, with sentence length RDM as a covariate. Bonferroni correction p< 0.05/7.

3. Control stage analysis

**The control verbal report stage.**

Fifty-one of the one hundred and one participants completed an additional control verbal report stage to map language processing areas (Fig. S7). This stage was a block design with two conditions. Participants were asked to look at the fixation cross on the screen, and when the “report” clue word appeared, they were instructed to report any thoughts or images that came to mind during the following time. If there were no thoughts/images in the participant’s mind at the stage where reporting was required, then they did not need to report. Participants were required to report only when thoughts/images were present during the reporting phase. When the clue word “don’t report” appeared, participants did not need to make any verbal report during the subsequent time. That is during the non-reporting phase, any thoughts/images that came to the participant’s mind did not need to be reported verbally. The duration of the cue representation was 1 second. It was worth noting that there was a 32-second buffer time before the start of the formal experiment, with a cross-fixed point presented on the screen. The sequence of reporting and non-reporting phases was presented in a pseudo-randomized form. Participants completed a total of 30 blocks, including 16 “report” blocks and 14 “don’t report” blocks. Each block lasted 20 seconds. In addition, each block was buffered by an empty screen. The buffered screens were presented in a pseudo-random form with a total time of 59.4 seconds. Therefore, the total scan time for the control verbal report stage was approximately 12 minutes. The order of the free verbal report stage and control verbal report stage was counterbalanced across participants. After quality control, the sample size for the data analysis of the control verbal report condition for language processing was 47.

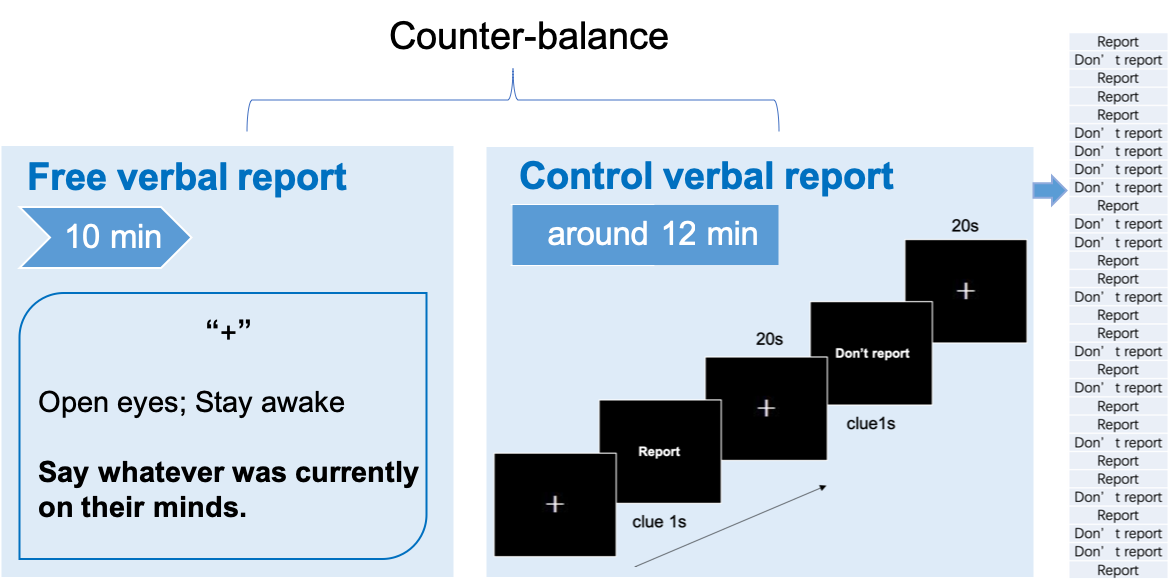

Fig. S7. Schematic diagram of the fMRI task. Fifty participants completed only the free verbal report stage, and fifty-one participants completed two stages. The first stage was the think-aloud paradigm which asked participants to free verbally report whatever came to their mind. The second stage was the control verbal report stage, where cue words indicate whether the participant needs to report or not. Clue words were presented in a pseudo-random form. The order of the latter two stages was counterbalanced across participants. The rightmost table shows the order in which the two blocks of the control verbal report stage are presented.

**The fMRI data preprocessing and analysis.**

The fMRI data preprocessing in the control condition was the same as for the free verbal report. SPM12 was used for individual-level analyses; a general linear model (GLM) was estimated for each voxel. The design matrix of the control verbal report stage was comprised of reporting blocks, non-reporting blocks and the six head motion parameters. The regressors were convolved with the SPM canonical hemodynamic response function. The high-pass filter was set at 128 s. After GLM estimation, the contrast images of the reported and non-reported blocks were calculated. Group-level analyses were assessed using whole-brain con-weight images. Multiple comparisons correction was performed using the Gaussian Random Field (GRF): voxel-level p < 0.001, cluster-level p < 0.05.

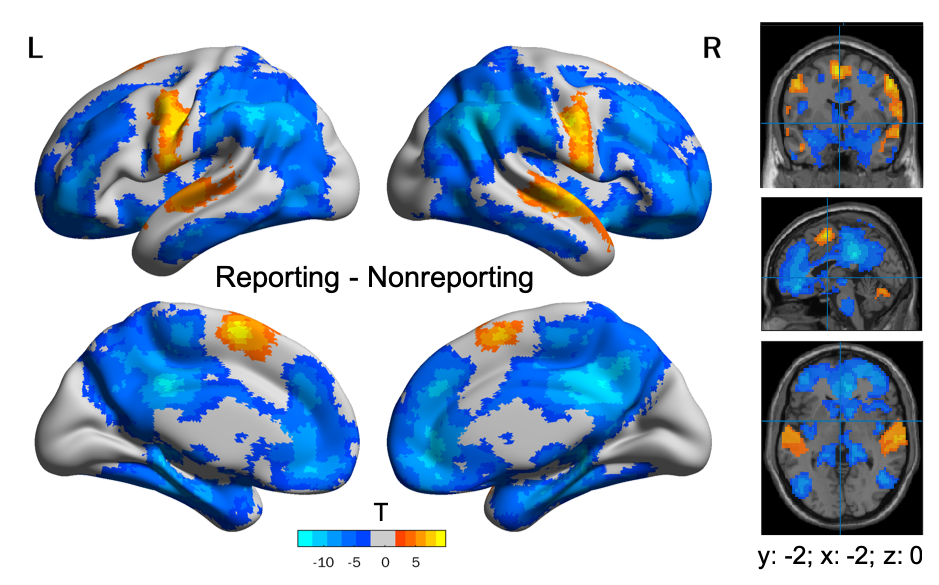

Fig. S8. The activation results of the reporting vs. non-reporting condition in the control report stage.

4. Other RSA results

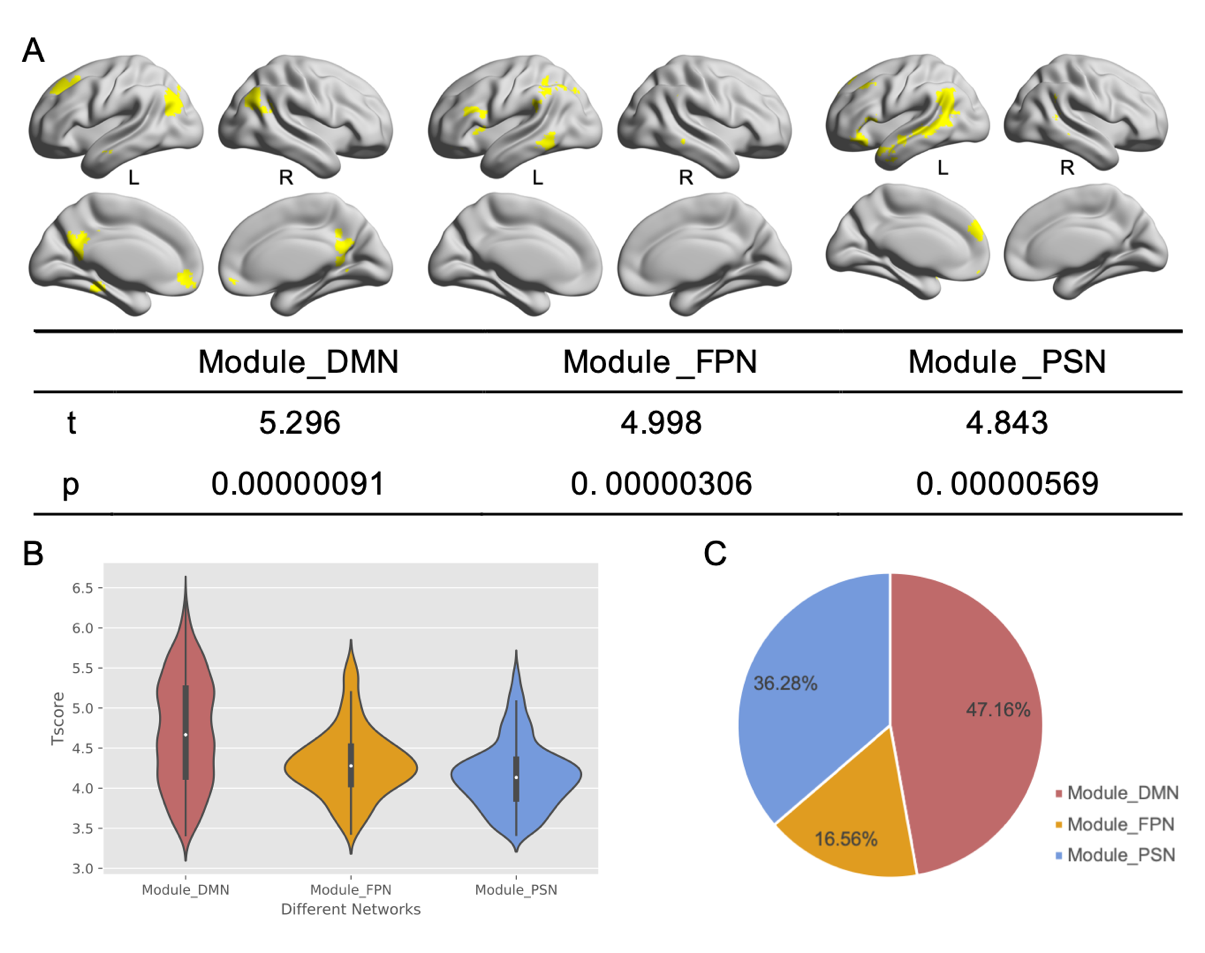

Fig. S9. A) RSA results were based on three semantically related modules. Sentence length RDMs were controlled as covariate. B) Distribution of significant voxels in three intrinsic functional semantic modules by whole-level searchlight RSA. C) Ratio of the number of significant voxels in the three semantic modules.

Table S4. RSA results for semantic RDM with random order based on the Yeo networks.

| Yeo seven-networks | t | p |
| --- | --- | --- |
| Visual network | -0.992 | 0.3239 |
| Somatomotor network | -1.500 | 0.1374 |
| Dorsal attention network | -0.524 | 0.6013 |
| Ventral attention network | -0.625 | 0.5339 |
| Limbic network | -0.864 | 0.3898 |
| Frontoparietal control network | -0.179 | 0.8586 |
| Default mode network | -0.797 | 0.4277 |

Note: We used the randomly disordered semantic RDM to compute the RSA.

5. Other related analysis results

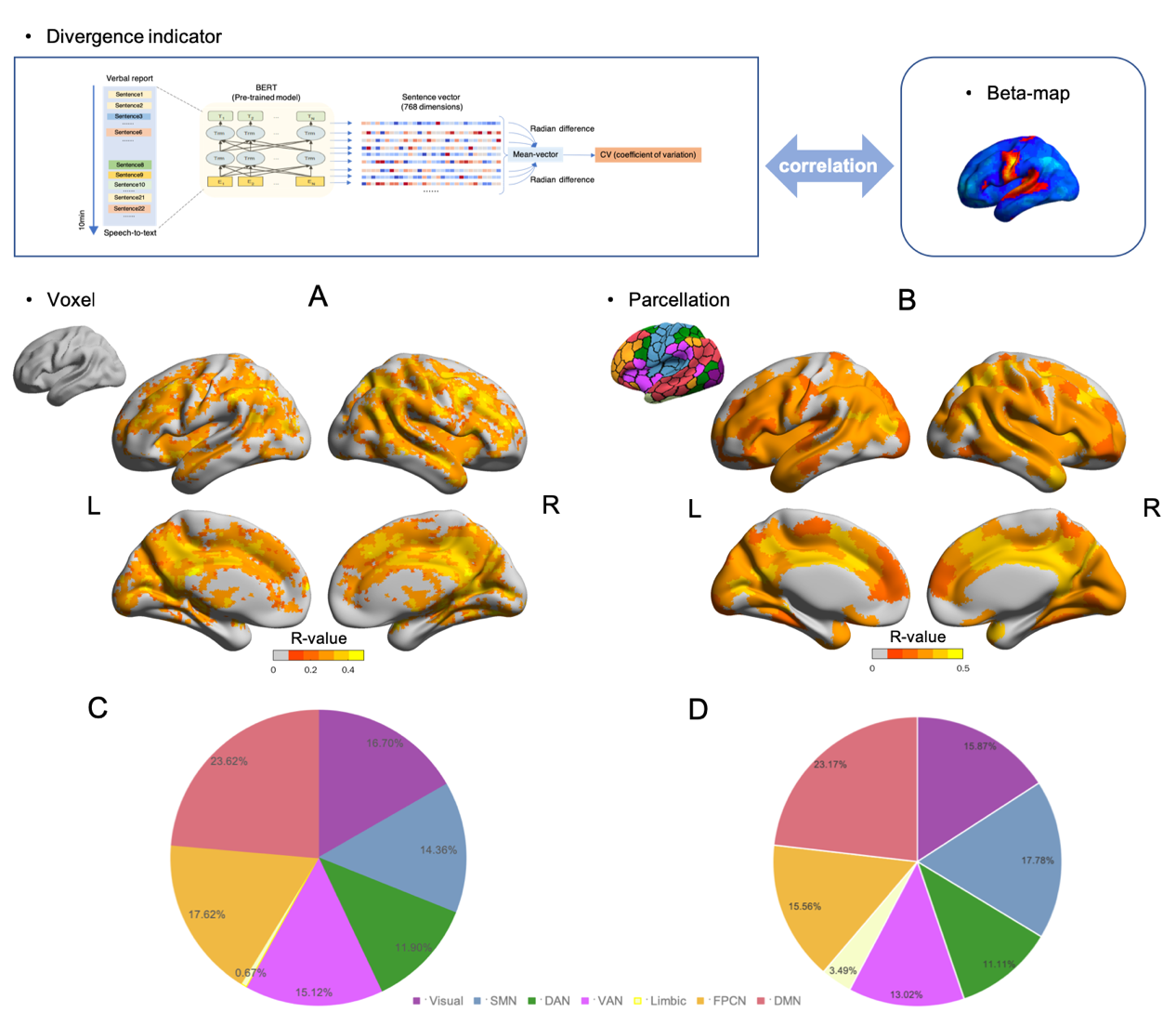

Fig. S10. Results of brain representations of fluctuations in thought content. Correlation results between the fluctuation indicator in thought content and the beta-map in the free verbal report stage based on voxels (A) and Schaefer 400-parcels (B). Multiple comparisons correction: Threshold-Free Cluster Enhancement (TFCE) with permutation test (Voxel-wise); FDR correction, q <0.05 (Schaefer 400-parcels). Significant voxels (C) and parcels (D) were distributed across seven Yeo networks. The ratio of the number of significant voxels/parcels divided by the total number of significant voxels/parcels.

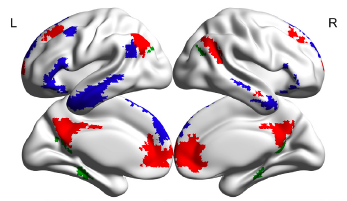

Fig. S11. The three subsystems of DMN were used in the present study. Red: core subsystem; Blue: DMPFC subsystem; Green: MTL subsystem.
